# Recessive antimorph alleles reveal novel functions of the OPAQUE1 myosin XI in maize

**DOI:** 10.1101/2025.06.26.661838

**Authors:** Brian Zebosi, Stephanie E. Martinez, Kokulapalan Wimalanathan, John Ssengo, Gabriela Brown, Norman B. Best, Michelle Facette, Carolyn G. Rasmussen, Erik Vollbrecht

**Affiliations:** Department of Genetics, Development and Cell Biology, Iowa State University, Ames, IA 50011; Interdepartmental Genetics and Genomics Graduate Program, Iowa State University, Ames, IA 50011; Botany and Plant Sciences Department, University of California, Riverside, Riverside, CA 92507; Interdepartmental Bioinformatics and Computational Biology Graduate Program, Iowa State University, Ames, IA 50011; Department of Plant Pathology, Iowa State University, Ames, IA 50011; USDA-ARS, Plant Genetics Research Unit, Columbia, MO 65211; Department of Biology, University of Massachusetts, Amherst, Amherst, MA 01003

## Abstract

Ideal plant architecture optimizes canopy structure and increases grain yield in maize. However, its underlying genetic mechanisms remain poorly characterized. Two recessive, EMS-induced maize mutants were identified that have reduced stature, opaque kernels, and abnormal subsidiary cell division, and are allelic to *opaque1 (o1)*. These two new missense alleles, *o1-2995* and *o1-tan62*, unlike the loss-of-function alleles previously identified, generate O1 protein but paradoxically generate more severe morphological defects. These defects include reduced internode elongation and partial suppression of excessive tassel and ear branching in *ramosa1 (ra1), ra2, and ra3* mutants. We show that *o1-2995* and *o1-tan62* are novel alleles of *o1*, and play a role in plant growth via internode elongation, subsidiary cell division positioning, leaf patterning, inflorescence development, and overall plant architecture.

## Introduction

Altering maize plant architecture traits has led to significant increases in yield <u>(Tian et al. 2019)</u>. Modern hybrids typically have narrow, erect leaves and tassels with few and upright branches that facilitate enhanced photosynthetic efficiency, efficient light interception, and tolerance to plant lodging under high planting densities <u>(Lambert and Johnson 1978; Tian et al. 2019;</u> <u>Pendleton et al. 1968)</u>. A key aspect of development in grasses is the regulation of branching in grain-producing inflorescences such as the tassel and ear of maize. Inflorescence branching architecture reflects the presence and differential activity of multiple axillary meristems. Prior research has identified several genes required for initiating axillary meristems and maintaining their determinacy. Some of these genes such as *Barren inflorescence1* (*Bif1*), *Bif2*, *Bif4*, and *Barren stalk 1* (*Ba1*) control auxiliary meristem initiation and their mutants display decreased tassel branching <u>(Barazesh and McSteen 2008; Gallavotti et al. 2004; Galli et al. 2015; McSteen</u> <u>et al. 2007; McSteen and Hake 2001)</u>. In contrast, the *ramosa* genes (*Ra1*, *Ra2*, and *Ra3*) regulate spikelet pair meristem (SPM) determinacy, and their loss of function mutants produce more tassel and ear branches <u>(Bortiri et al. 2006; Satoh-Nagasawa et al. 2006; Vollbrecht et al.</u> <u>2005)</u>. *Ra1* encodes a C2H2-type zinc finger transcriptional regulator that likely acts as both an activator and repressor of target genes <u>(Gallavotti et al. 2010; Eveland et al. 2014; Vollbrecht et</u> <u>al. 2005)</u>. *Ra2* encodes a LATERAL ORGAN BOUNDARIES domain transcription factor <u>(Bortiri et al. 2006)</u>. *Ra3* encodes a trehalose-6-phosphate phosphatase whose role in branching is uncoupled from enzymatic activity <u>(Claeys et al. 2019; Satoh-Nagasawa et al. 2006)</u>). Using a genetic approach to identify modifiers of *ramosa1* (*ra1*), we uncovered a novel mutant that suppresses tassel branching independent of the *ramosa* pathway and that disrupts a gene encoding a myosin XI-I called *Opaque1* <u>(Wang et al. 2012)</u>.

Myosins are molecular motors that transport cargo along cytoskeletal actin filaments (F-actin) and are required for proper cell function. Myosins are important for processes such as organelle movement, cell shape maintenance, cell polarization, actin organization and signal transduction <u>(Madison and Nebenführ 2013)</u>. The myosin protein structure typically consists of three domains: the head, neck, and tail <u>(Foth et al. 2006; Tominaga and Nakano 2012)</u>. The head motor domain functions in actin binding and in ATP binding and hydrolysis for motor activity, the neck domain contains a linker that influences motor step size, while the tail binds cargo and facilitates dimerization <u>(Nebenführ and Dixit 2018)</u>. Myosins are highly conserved across many eukaryotic species, including mammals, fungi, and plants <u>(Richards and Cavalier-Smith 2005)</u>. Two distinct myosin groups (VIII and XI) are specific to plants and present as several subfamilies due to repeated rounds of whole-genome duplications <u>(Foth et al. 2006; Mühlhausen</u> <u>and Kollmar 2013; Nebenführ and Dixit 2018)</u>. Genetic studies and functional characterization of mutants of these plant-specific myosins indicate they are involved in many processes including organelle translocation, cytoplasmic streaming, cytokinesis, tip growth, exocytosis and endocytosis <u>(Nebenführ and Dixit 2018; Zhang et al. 2019; Chocano-Coralla and Vidali 2024;</u> <u>Golomb et al. 2008; Peremyslov et al. 2010; Olatunji et al. 2023; Tominaga and Nakano 2012;</u> <u>Ueda et al. 2015)</u>.

In particular, Arabidopsis myosin XI-I isoforms (myosin XI-Is) are localized to the nuclear envelope and act as linkers between the nucleus and actin cytoskeleton through a linker of nucleoskeleton and cytoskeleton (LINC) complex <u>(Tamura et al. 2013)</u>. Arabidopsis myosin XI-I mutants (*Atxi-1*/*kaku1*) have aberrant nuclear envelopes with impaired nuclear positioning and movement, thus emphasizing the vital role of myosins in nuclear movement and shape maintenance <u>(Tamura et al. 2013; Muroyama et al. 2020)</u>. Although important for nuclear movement in Arabidopsis, genetic studies in maize indicate that myosin XI-I is instead required for protein body formation during seed development and endoplasmic reticulum (ER) movement <u>(Wang et al. 2012)</u>. Additionally, the *opaque1* mutants have phragmoplast guidance defects leading to asymmetric cell division defects <u>(Nan et al. 2023)</u>. Unlike *kaku1* mutants, no nuclear movement abnormalities were observed in *o1* asymmetric divisions <u>(Nan et al. 2023)</u>. However, neither growth nor developmental defects were reported in these myosin XI mutants.

Here, we functionally characterized two new alleles of *opaque1,* a myosin XI-I (Zm00001eb193160). One mutant allele, *ramosa1 suppressor locus*12.2995* (*rsl*-12.2995*), was identified from a *ramosa1* ethyl methanesulfonate (EMS) enhancer-suppressor screen as a mutant with reduced stature, abnormal subsidiary cell division, opaque kernels and impaired tassel, ear and vegetative shoot development. The *rsl*-12.2995* mutation was mapped to the *O1* locus using map-based cloning and whole-genome sequencing and subsequently named *o1-2995*. The other mutant allele, *tangled62* (*o1-tan62*), was identified from a forward genetics EMS screen for plants with abnormal subsidiary cell divisions. Allelism tests confirmed both as new *o1* alleles that in genetic analyses behaved as recessive antimorphs. Thus, *o1-2995* and *o1-tan62* are novel alleles of *Opaque1 (O1)* that regulate plant growth, architecture and asymmetric division positioning in maize.

## Results

### The *o1-2955* mutant shows aberrant shoot and inflorescence architecture

Genetic and genomics studies indicate that three *Ramosa* genes in maize (*Ra1, Ra2* and *Ra3*) regulate branching in grass inflorescences by modulating shared and discrete developmental modules <u>(Vollbrecht et al. 2005; Bortiri et al. 2006; Satoh-Nagasawa et al. 2006; Eveland et al.</u> <u>2014)</u>. To identify *ramosa1* (*ra1*) mutant modifiers, we performed an EMS mutagenesis screen of *ra1-63* mutants <u>(Vollbrecht et al. 2005)</u> in the Mo17 genetic background. In the M2 generation, we identified a short-statured mutant that suppressed *ra1*-induced tassel and ear branching, named it *ramosa suppressor locus*-12.2995* (*rsl*-12.2995*) and later renamed it *o1-2995*. In F2 and backcross populations, *o1-2995* segregated as a single recessive locus (wildtype:*o1-2995*, 48:22 [χ2, P = 0.21] and 22:18 [χ2, P = 0.53], respectively). *o1-2995* single mutants also showed reduced stature after backcrossing 5-8 times into B73 and W22 (**Figure 1D and 1E**). The *o1-2995* mutant’s effect on plant height was completely penetrant in all three genetic backgrounds, but its expressivity varied. Compared to their normal siblings, field-grown *o1-2995* mutants were shortest in B73, suggesting that genetic modifiers interact with *O1* in its effect on plant height. All subsequent phenotyping was carried out using the well-introgressed B73 stocks.

**Figure 1.**
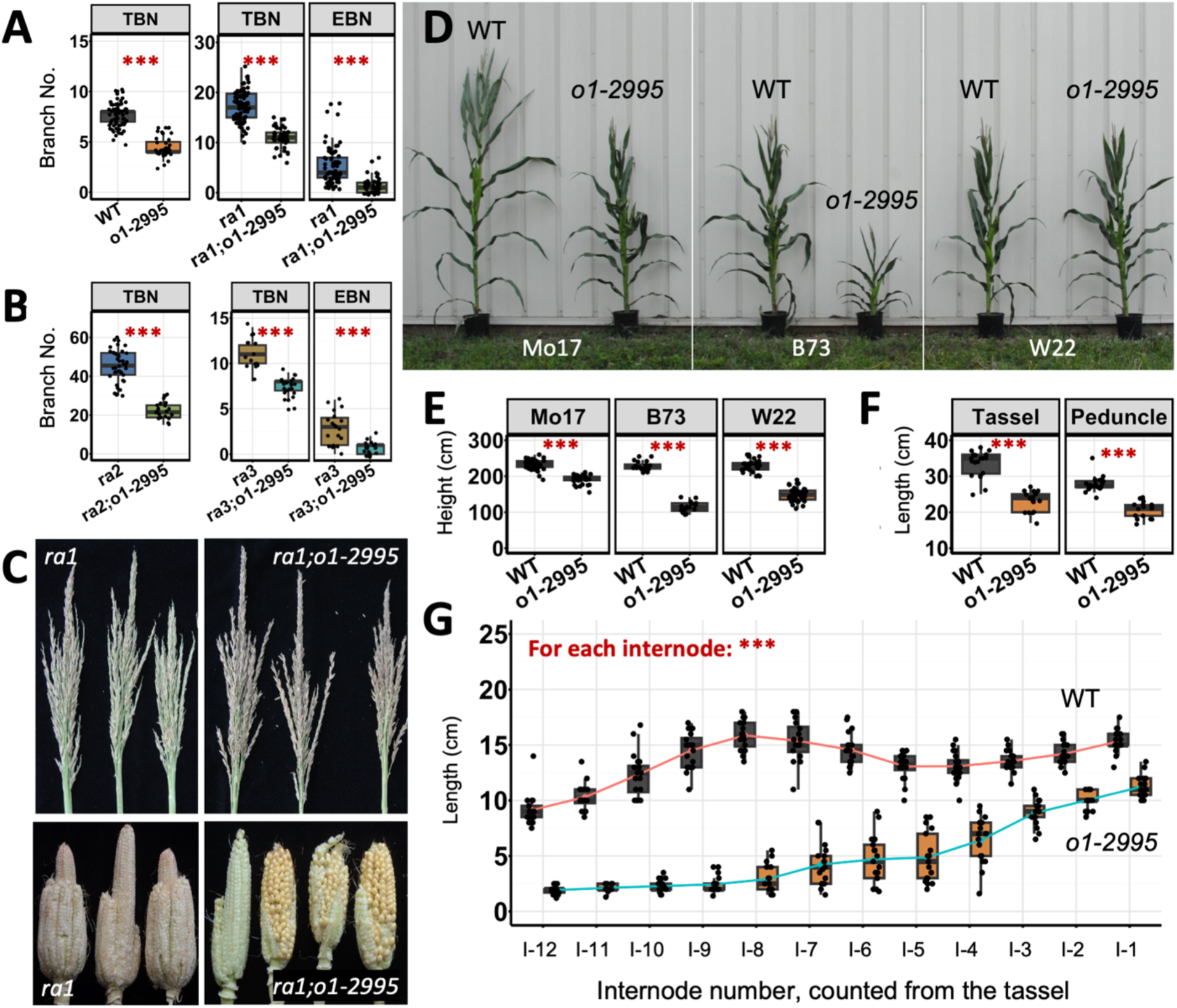
Phenotypic characterization of *o1-2995* and of its interaction with *ramosa* (*ra*) pathway genes. **(A-C)**. *o1-2995* and its interaction with *ra1*, *ra2* and *ra3* in B73. **(A).** Left, tassel branch number (TBN) of normal siblings (WT, n = 75) and *o1-2995* single mutants (n = 33). Right, TBN and Ear branch number (EBN) of *ra1* single mutants and *ra1;o1-2995* double mutants. **(B).** Left, TBN of *ra2* single mutants and *ra2;o1-2995* double mutants. Right, TBN and EBN of *ra3* single mutants and *ra3;o1-2995* double mutants. **(C)**. Representative tassels and ears as quantified in Panel A, right. **(D-E).** Plant height across inbred line (B73, Mo17 and W22) introgressions for normal siblings (WT, n = 47, 20, 21 for Mo17, B73, W22 respectively) and *o1-2995* mutants (n = 39, 17, 39 for Mo17, B73, W22 respectively). **(F)**. Peduncle length and whole tassel length of normal siblings (WT, n = 30) and *o1-2995* mutants (n = 30) in B73. **(G)**. Length of several internodes of the main shoot in normal siblings (WT, black, left, successive means connected by red line, n = 20) and *o1-2995* mutants (orange, right, successive means connected by blue line, n = 17) in B73. I-1 denotes the internode nearest the tassel and I-12 is 11 internodes away basally, near the soil line. Asterisks (***), significant difference at P < 0.001.

In B73, tassel branch number was reduced in the *o1-2995* single mutant which produced 51.1% fewer tassel branches than its wildtype siblings (**Figure 1A**). Length of the tassel peduncle and tassel length exhibited a 70% and 78.1% reduction, respectively, in mutants compared to wildtype siblings (**Figure 1F**).

Because *o1-2995* was isolated from a *ra1-63* mutant suppressor genetic screen, we constructed double mutants between *o1-2995* and all three *ramosa* pathway mutants (*ra1*, *ra2*, and *ra3*), and also produced one triple mutant (*ra1;ra2;o1-2995*). We compared the higher order mutant phenotypes to the single mutants to determine if *O1* and *Ra* genes function in an additive, synergistic, or epistatic manner with one another. In *ra1*, *ra2* or *ra3* mutants, additional long branches form in both the ear and in the tassel. As stated earlier, the *o1-2995* single mutant reduced tassel branch number (TBN) by 51.1% as compared to its wildtype siblings (**Figure 1A, left; 1C**). *o1-2995* single mutants did not reduce ear branch number (EBN) as the normal ear is unbranched. In double mutant populations with *ra1-63,* the *o1-2995* mutant reduced TBN by 36.5% and EBN by 72.3% (P < 0.001, Student’s t-test), as compared to *ra1-63* single mutants (**Figure 1A, right; 1C**). Thus, in combination with *ra1-63*, the *o1-2995* mutant behaved additively by slightly reducing overall branching. The *ra2-R;o1-2995* double mutant combination also appeared additive with a 51.4% reduction in TBN in the double mutant as compared to the *ra2-R* single mutant (**Figure 1B, left**). In mutant combinations with *ra3*, *o1-2995* also additively suppressed tassel and ear branching; the *ra3-R*;*o1-2995* double mutants exhibited 33.4% and 76.6% reductions in TBN and EBN, respectively, compared to *ra3-R* single mutants (**Figure 1B, right**). Thus, for all three double mutants (*ra1-63*;*o1-2995*, *ra2-R*;*o1-2995* and *ra3-R*;*o1-2995*) the inflorescence branching phenotypes provided no clear evidence of epistasis, as if *o1-2995* affected growth more generally without specifically affecting the meristem determinacy regulated by the *ramosa* genes. As a final test we examined *ra1;ra2* double mutants which interact synergistically by more completely reducing activity of the *ramosa* pathway, leading to profusely branched ears <u>(Vollbrecht et al. 2005)</u>. The (*ra1-63;ra2-R;o1-2995*) triple mutants had substantially less ear branching and smooth patches on the ear axis, which we interpreted as additive or possibly synergistic, since the patches are not seen in either single or double mutant (**Supplementary** Figure 1B). This implies that *O1* and *ramosa* genes are in either independent or converging genetic pathways, and are unlikely to participate together in a simple linear pathway to regulate inflorescence branching.

### Map-based cloning and identification of *o1-2995* as a novel allele of *Opaque1*

To identify the causative mutation responsible for the *rsl*-12.2995* (*o1-2995*) mutant phenotype, we used map-based cloning methods. Bulked segregant analysis using genotyping-by-sequencing (BSA-GBS) mapped the *o1-2995* mutation to the long arm of chromosome 4 (**Supplementary** Figure 2), within a peak containing 989 genes. Mapping using publicly available molecular markers identified a 1.5Mb interval containing 28 genes (**Figure 2A**) which, in non-repetitive regions, contained no DNA sequence polymorphisms between B73 and Mo17. We therefore performed whole-genome sequencing (WGS) of a single mutant plant and identified EMS-induced SNPs in the interval. Fine-mapping in a second, larger (2289 plants) population indicated that the causal mutation was within a physical distance of ∼ 0.6Mb that contained only one gene, the previously cloned *Opaque1* (*O1*) (**Figure 2B**). We therefore phenotyped kernels from our segregating populations, and found an opaque kernel phenotype co-segregated with the *rsl*-12.2995* mutant plant phenotypes.

**Figure 2.**
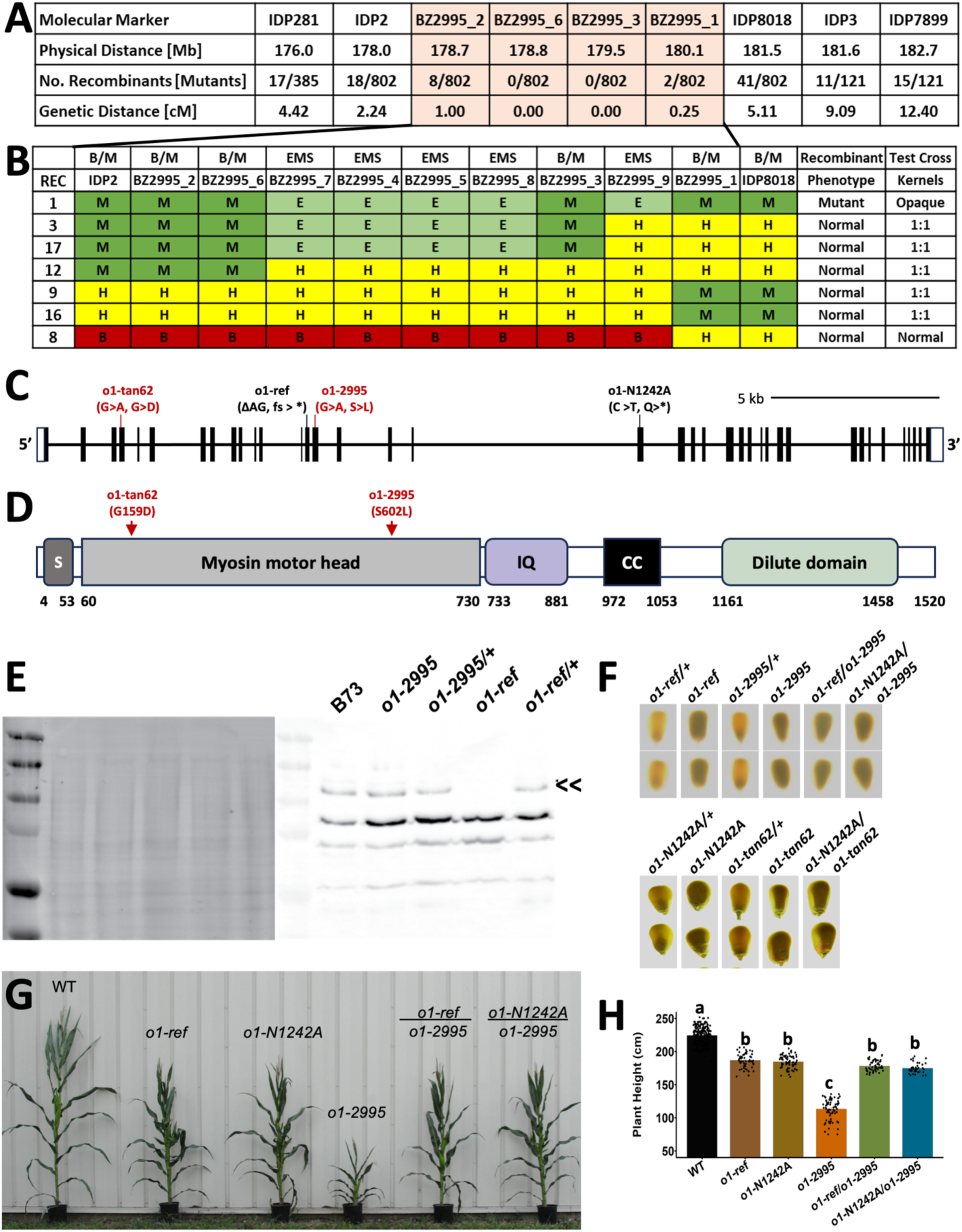
Map-based cloning, *o1-2995* and *o1-tan62* complementation tests and antimorph allele relationships. **(A-B)**. Map based cloning of *o1-2995*. **(A)**. Highlighted interval (pink) is ∼1.5MB with 28 genes and no B73/Mo17 polymorphism. **(B).** Whole-genome sequencing-aided recombination mapping. E, EMS-like SNP. M, homozygous Mo17. H, heterozygous Mo17/B73. B, homozygous B73. The new interval contained eight genes and only one exonic, EMS-like SNP (G>A), in the *o1* gene. **(C).** *o1* intron-exon gene structure (from NCBI) showing relative mutation sites of *o1-2995* and *o1-tan62* and of null alleles *o1-N1242A* and *o1-ref* (Wang et al. 2012) used in this study. **(D)** Maize O1 protein domain structure and relative locations of *o1-2995* and *o1-tan62* missense mutations in the myosin head domain. S, SH3-like domain; IQ, region of six IQ domains; CC, coiled-coil domain. **(E)**. SDS-PAGE loading control (left) and anti-O1 western blot (right) of membrane and membrane-associated proteins isolated from the stomatal division zone of wild type (B73) and *o1* mutant plants. Double arrowhead, migration position of the O1 protein. **(F, G and H)**. Complementation test for plant height and kernel phenotypes. **(F)**. Kernels of indicated genotypes, illuminated from below on a light box; *o1-2995* and *o1-tan62* fail to complement endosperm phenotype of null alleles *o1-ref* and *o1-N1242A*. **(G and H)**. Field-grown plants of indicated genotypes in B73; *o1-2995* fails to complement plant height phenotype of null alleles *o1-ref* and *o1-N1242A*. Panel H, for the genotypes left to right, n = 70, 29, 39, 36, 59, 37, respectively; letters a, b, c, significant groupings by ANOVA with Tukey’s HSD.

Sanger sequencing of the *O1* locus in *o1-2995* confirmed the single nucleotide change (G > A) in the 15^th^ exon (**Figure 2C**), causing a missense mutation (S602L) within the myosin head domain of O1 (**Figure 2D**). S602 is a nearly-invariant residue near the actin-binding motif: across the 72 myosin VIII and myosin XI genes from flowering and non-flowering plants we surveyed for Figure 3, 71 genes encode an S in this position and one encodes a T.

**Figure 3.**
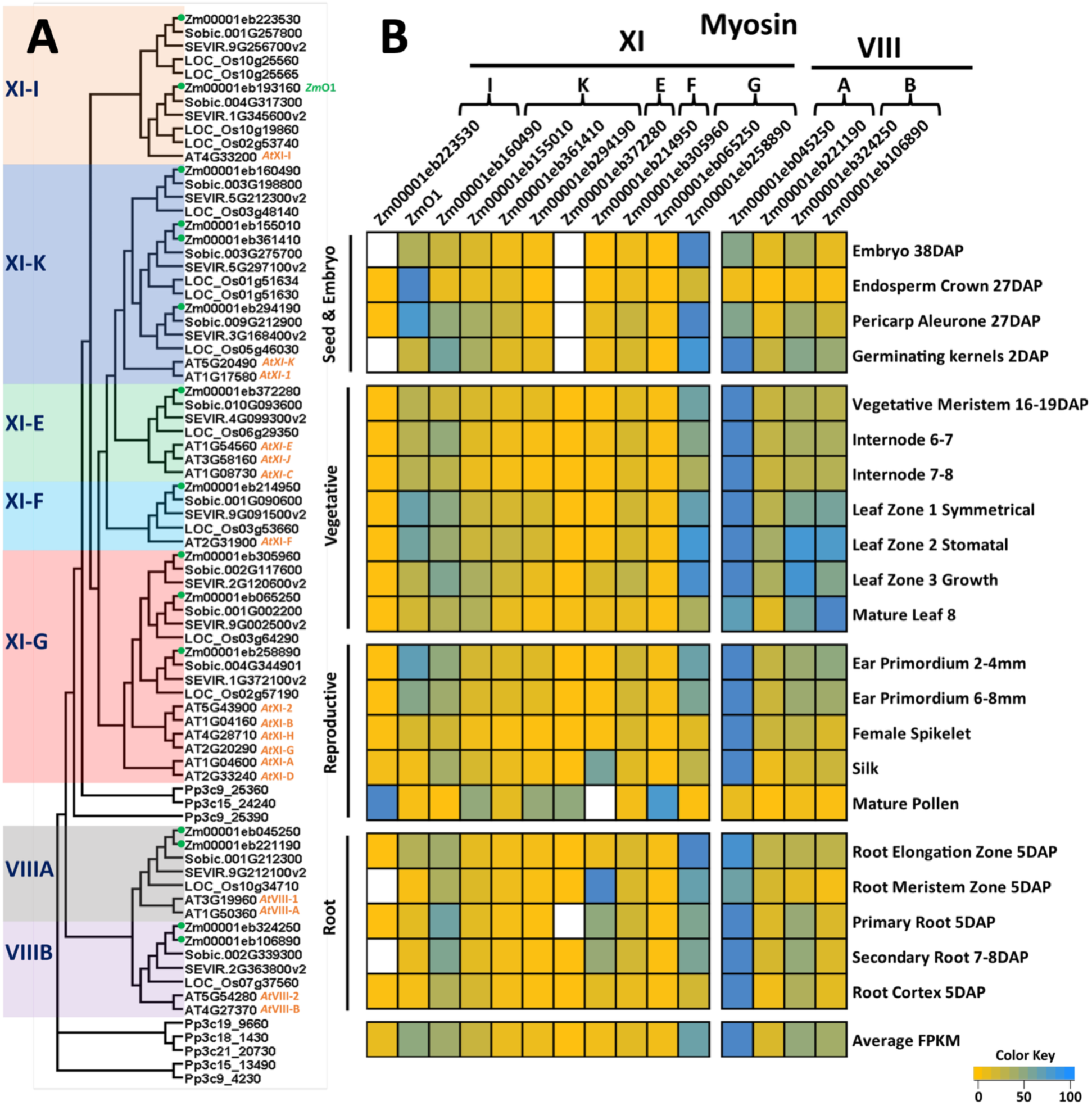
Phylogenetic and gene expression analysis of plant myosins XI and VIII. **(A)**. Phylogenetic analysis of all plant myosins XI and VIII across several species. Zm, maize; Sobic., *Sorghum bicolor*; SEVIR, *Seteria viridis*; Os, rice; AT, Arabidopsis; Pp, *Physcomitrella patens* (moss) as an outgroup. (**B).** Relative transcript expression levels of myosin XI and VIII genes in different maize tissues; from Walley et al., (2016).

To confirm the genetically linked, EMS-induced mutation in *O1* as causal of the *o1-2995* phenotype, we performed complementation tests with previously identified null mutants, *o1-ref* and *o1-N1242A* <u>(Wang et al. 2012)</u>. *o1-2995* failed to complement the *o1* null alleles for kernel opaqueness (**Figure 2F**). Interestingly, we also discovered that each null mutant, *o1-ref* and *o1-N1242A*, showed slightly reduced plant height (**Figure 2G and 2H**), a previously unreported phenotype. Moreover, the reduced plant height of the compound heterozygous mutants (*o1-ref*/*o1-2995* and *o1-N1242A*/*o1-2995*) was similar to *o1-ref* or *o1-N1242A* homozygotes. Plant height was greatly reduced for *o1-2995* mutants (**Figure 2G and 2H; see also Figure 4A**). Thus, the *o1-2995* allele generates a stronger mutant phenotype when homozygous than do null mutant alleles *o1-ref* or *o1-N1242A*. These data imply that *o1-2995* functions as a formally recessive but dose-dependent, antimorphic, allele of *O1*.

**Figure 4.**
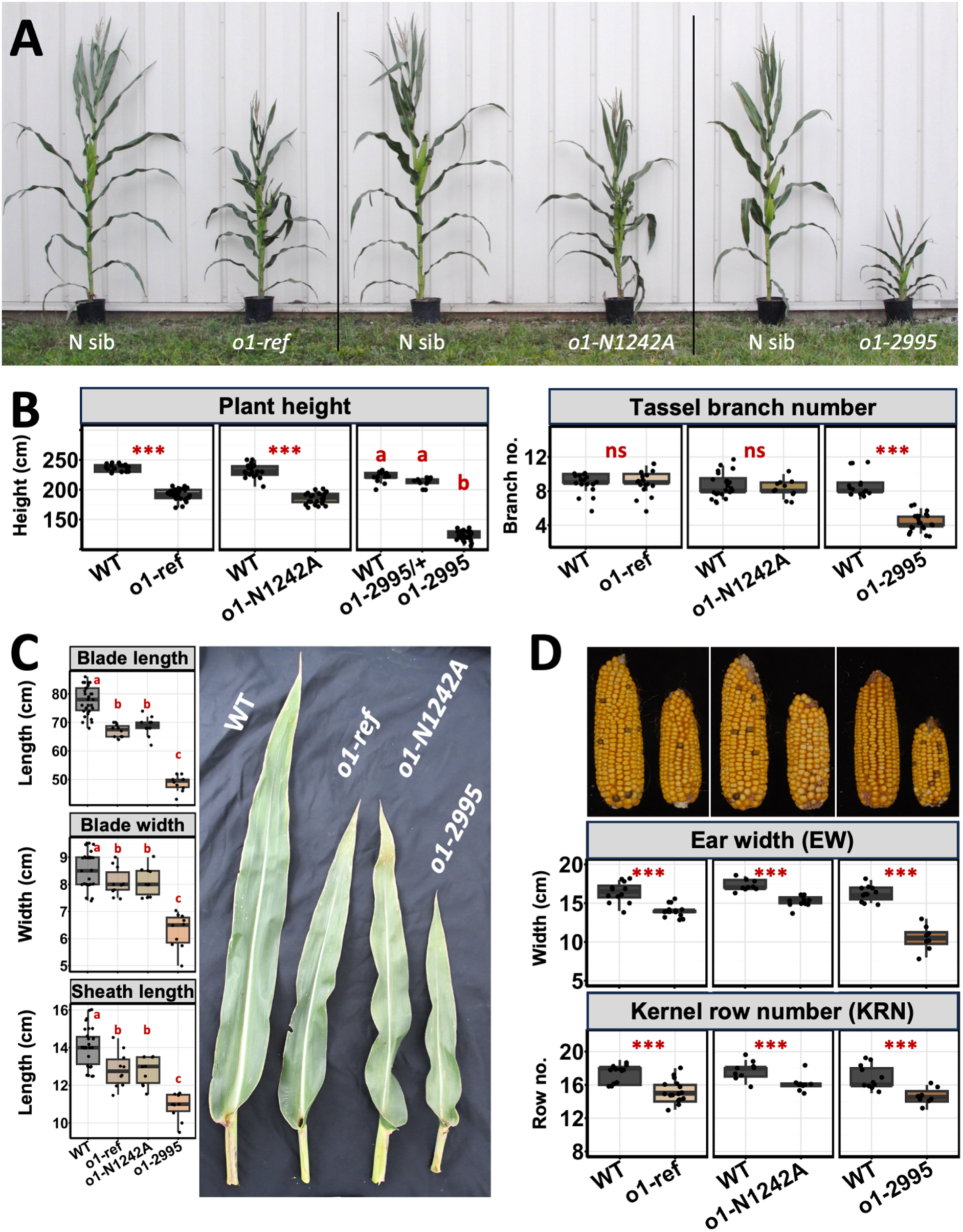
Shoot phenotypes of *o1* mutant alleles in B73. **(A)**. Representative field-grown plants of indicated genotypes and their wild type (N) siblings. **(B)**. Plant height (left) and tassel branch number (right) of indicated genotypes and wild type (WT) siblings. **(C)**. Quantification of leaf characters (left) and images of representative leaves (right) of WT (B73) and indicated mutant genotypes. **(D)**. Ears (top) and quantification of ear size characters of indicated genotypes and wild type (WT) siblings. Error bars, standard error; asterisks (***), significance at P < 0.001 by t-test; letters a, b, c, significant groupings by ANOVA with Tukey’s HSD; ns, no significant difference.

To investigate if the non-synonymous point mutation in *o1-2995* altered O1 protein levels, we isolated proteins from the stomatal division zone of the leaf and used an O1-specific antibody to compare amounts of protein produced between *o1-ref* and *o1-2995*. As expected, a band was detected in the normal inbred B73 plants and wildtype, heterozygous siblings of *o1-ref* and *o1-2995*, while no band corresponding to the size of the O1 protein was detected in *o1-ref* mutant plants (**Figure 2I**). In contrast, in *o1-2995* mutants, a band of the expected size was detected, indicating the mutant produces a protein (**Figure 2I**). These data indicate that *o1-2995*, which expresses myosin XI-I containing a S602L mutation in the motor domain, confers a stronger mutant phenotype than *o1* null mutants which express no or very little myosin XI-I protein.

### *o1-tangled62* is another novel allele of *opaque1*

An additional mutant allele of *o1* was identified through a forward genetics screen to identify mutants with division plane orientation defects. A recessive mutant with aberrant subsidiary cells was initially identified as *tangled62* (*tan62*) (**Supplementary** Figure 3). Due to the similarity of the subsidiary cell division defect compared to *o1* mutants as well as an opaque kernel phenotype <u>(Nan et al. 2023; Wang et al. 2012)</u>, complementation tests were performed with *o1-N1242A* mutants. Across 11 independent crosses, *tan62* failed to complement the *o1-N1242A* opaque kernel phenotype, suggesting that *tan62* is allelic to *o1* (**Figure 2F**). Sequencing of the *tan62* cDNA and DNA extracted from three additional *tan62* mutants revealed a single nucleotide change (G > A) in the 4^th^ exon (**Figure 2C**), causing a missense mutation (G159D) within the myosin motor head domain (**Figure 2D**). G159 is an invariant residue in the head domain of all myosins VIII and XI across flowering and non-flowering plants surveyed in Figure 3, located in the ATP binding pocket. This type of mutation is predicted to be deleterious but still produce protein, similar to O1-2995. Therefore, *tan62* was renamed to *o1-tan62*.

### Genome-wide identification and functional analysis of myosin XIs in maize

To consider novel *O1* alleles in the context of the genome-wide diversification of myosins in maize and other plants, we constructed a phylogenetic tree and annotated it with expression data using publicly available sources <u>(Walley et al. 2016; Wang et al. 2012)</u>. Using blast searches based on Arabidopsis myosin XI and VIII protein sequences, we included all relevant amino acid sequences from Arabidopsis, rice, sorghum, *Setaria* and as an out-group, *Physcomitrium patens* (moss) (**Figure 3A**). Within the myosin XI clade, Arabidopsis, maize and rice have 13, 12 and 11 genes, respectively, while Setaria and sorghum have 10 myosin XIs. In general (**Figure 3A**), the myosin XIs are further clustered into five distinct subclades ( XI-I, XI-K, XI-E, XI-F, and XI-G) based on the Arabidopsis nomenclature <u>(Avisar et al. 2008; Reddy and Day 2001;</u> <u>Peremyslov et al. 2011)</u>. *Opaque1* (*O1*) belongs to the myosin XI-I clade, which comprises two genes in maize, Setaria and sorghum, contrasting to one in Arabidopsis and four in rice. Spatial expression differences among the different maize myosins (**Figure 3B**) were annotated from publically available expression data <u>(Walley et al. 2016; Wang et al. 2012)</u>. The two myosin XI-I genes in maize (Zm00001eb223530 and *O1*) exhibited opposite expression patterns. Notably, *O1* is expressed more broadly and more highly than Zm00001eb223530 in most tissues surveyed except mature pollen, which implies functional diversification between the two genes and suggests they function non-redundantly.

### Effects of different *o1* mutant alleles on plant shoot phenotypes and in different environmental conditions

*opaque1* has been studied for over 50 years as a classical endosperm mutant <u>(Wang et al. 2012;</u> <u>Neuffer et al. 1968)</u>yet only recently have any plant shoot phenotypes been reported in *o1* loss of function mutants, for a role in asymmetric cell divisions in the leaf <u>(Nan et al. 2023)</u>. Because *o1-2995* antimorph allele mutants had altered shoot development phenotypes such as tassel branching and plant height (**Figure 1**) and both *o1-2995* and *o1-tan62* failed to complement the endosperm phenotype of *o1* null alleles (**Figure 2F**), we hypothesized that additional shoot phenotypes would be affected by various alleles, and tested this hypothesis using our mutant stocks in the B73 genetic background.

### Field-grown plant phenotypes indicate a role for *O1* in plant height and ear growth

In side-by-side field growth experiments comparing single mutants of the antimorphic *o1-2995* allele and the null alleles *o1-ref* and *o1-N1242A*, the *o1-2995* plants were again dramatically smaller (45.8% height reduction) than *o1-2995/+* or homozygous +/+ siblings. The *o1-ref* and *o1-N1242A* mutants also showed height reduction compared to their wild type siblings, although the reduction was less severe than in *o1-2995* (18.7% and 19.6% reduction, respectively) (**Figure 4A, 4B**). On the other hand, *o1-2995* mutants had 3-5 fewer tassel branches but *o1-ref* and *o1-N1242A* null mutant alleles had no significant effects on TBN relative to their wildtype siblings (**Figure 4B, right**). Finally, the length and width of ears, and kernel row number, was significantly reduced in all mutants compared to their wild type siblings, with more abnormal phenotypes observed in *o1-2995* mutants than in the null allele mutants (**Figure 4D**). These data suggest that *O1* plays a greater role in plant height, and ear growth, than tassel branching.

### Greenhouse-grown plant phenotypes suggest *O1* is responsive to environmental conditions

Greenhouse growth experiments also showed reductions in plant height in some of *o1* mutants compared to wild-type siblings, but with more variability. When *o1-2995* mutants were grown in the greenhouse during the shorter days of winter 2018, mutants were significantly shorter than their wildtype siblings (average 179.7 cm and 225.8cm, respectively; P = 0.00042), although proportionally less so (20.2%,) compared to the reduction in field-grown plants (45.8%). In a separate greenhouse experiment performed in 2024, both *o1-2995* and *o1-tan62* mutants had a significant reduction in plant height of 37.8% and 17.1%, respectively, compared to wild-type siblings, while *o1-N1242A* null mutants did not, when measured at the whole plant level (**Supplementary** Figure 5A). Similarly, peduncle length, tassel length and TBN were unaffected in *o1-N1242A* but were generally reduced in *o1-tan62* and *o1-2995* mutants. Thus, environmental conditions may modulate the severity of phenotypic effects but under all conditions tested *o1-2995* and *o1-tan62* had more reduced stature compared to both wild-type siblings and to null mutants *o1-ref* and *o1-N1242A*.

### Plant height reduction is due to compressed internodes

To understand the basis of reduced stature of the *o1-2995* mutants, we counted the number of leaves formed and measured the size of internodes. Our results showed that the normal siblings and mutants generated similar numbers of leaves (normal, n=100, 20.2 leaves; *o1-2995*, n= 42, 19.1 leaves; P=0.072) but that in mutants, internode lengths and diameters were compressed, with a 26-83% internode length reduction (P<0.0001) compared to wild-type siblings depending on the observed internode (**Figure 1G and Supplementary** Figure 4). The reduction in internode length varied but was more pronounced in the lower internodes (with 78.9% and 83.2% reduction) than in the internodes above the ear and towards the tassel. Similarly reduced internode lengths were also observed in *o1-tan62* mutants (**Supplementary** Figure 5B). Thus, the plant height reduction in the *o1-2995* and *o1-tan62* mutants is due to impaired internode elongation.

### *O1* functions in leaf cell expansion and leaf development

In side-by-side field growth experiments comparing single mutants of the antimorphic *o1-2995* allele and the null alleles *o1-ref* and *o1-N1242A,* we observed generally smaller and narrower leaves in all mutants as compared to their normal siblings. Mirroring the trend observed for plant height, length of leaf blades and sheaths, and width of leaf blades were slightly reduced in the null mutants, and more so in *o1-2995* antimorph mutants (**Figure 4C**). To determine the basis of small leaf size we examined *o1-2995* mutants by making abaxial leaf epidermal impressions from leaf 16 using super-glue and observed the impressions under a compound microscope. Compared to their wildtype siblings, *o1-2995* mutants had significantly more cells per microscope field of view (**Supplementary** Figure 6D) due to reduced cell size (**Supplementary** Figure 6C). Thus, the *o1-2995* mutation impairs cell expansion and growth.

In addition, *o1-2995* mutants displayed aberrant leaf patterning. Especially when grown in the field, *o1-2995* mutants had ectopic tissue outgrowths extending from the ligule into the leaf midrib (**Supplementary** Figure 7A-7C). To further investigate if *o1-2995* mutation also affected vein development, we examined acidified phloroglucinol stained cross sections of distal midribs of adult leaf 10 under a dissecting microscope. Mutants had markedly reduced volume of clear cells, the tissue located adaxial to the midvein that comprises the bulk of the midrib, accordingly reduced midribs, and smaller veins compared to their wild type siblings (**Supplementary** Figure 7D-7G). The ectopic outgrowths were not observed in *o1-2995* or *o1-tan62* mutants grown in the greenhouse, or in any *o1* null mutants grown under any conditions. These data indicate that the antimorph *o1-2995* allele disrupts ligule and midrib patterning and development, and does so more severely under field grown conditions.

### *o1* antimorph alleles show severe subsidiary cell division defects

*o1* null mutant alleles (*o1-ref* and *o1-N1243*) were reported to cause kernel opaqueness, impaired endoplasmic reticulum (ER) motility, irregular protein body shape, and aberrant stomatal subsidiary cell division (Nan et al., 2023; Wang et al., 2012). We showed previously that *o1* null mutants have abnormal subsidiary cells and defective phragmoplast guidance (Nan et al., 2023). From epidermal glue impressions of leaf 4 the *o1-ref* allele had 20% abnormal subsidiary cells, similar to previously published results, (Nan et al., 2023), while plants homozygous for the antimorph *o1-2995* allele had ∼45% abnormal subsidiary cells (**Figure 5A to 5F**). *o1-N1242A* mutants had ∼20% abnormal subsidiary cells (n=11 plants), also similar to previous reports <u>(Nan</u> <u>et al. 2023)</u>, while *o1-tan62* mutants had ∼30% abnormal subsidiary cells (n=15 plants) in juvenile leaf 2 (**Supplementary** Figure 3A). Thus, both antimorph alleles *o1-2995* and *o1-tan62* more severely disrupt cell divisions for subsidiary cells than *o1* null mutants.

**Figure 5:**
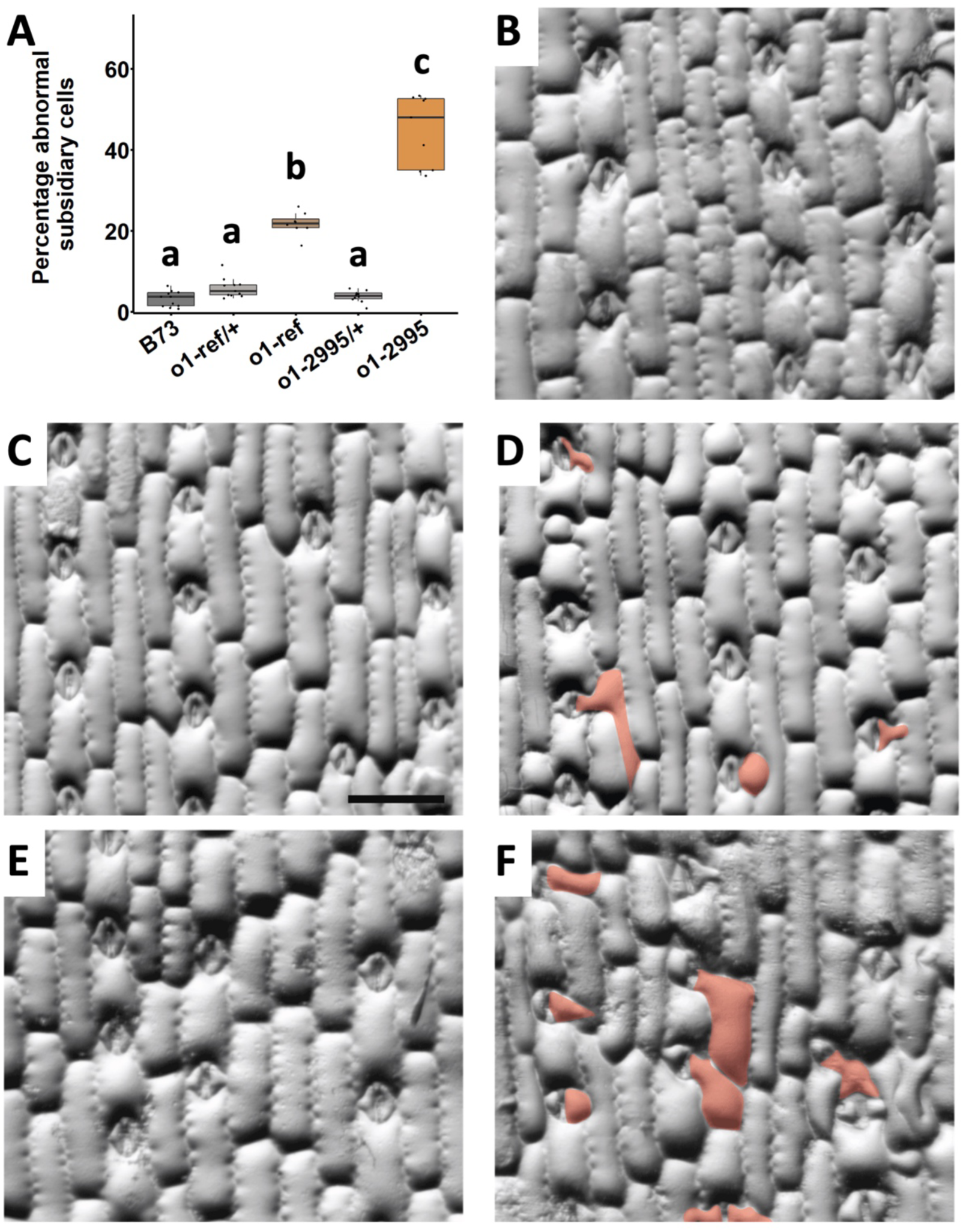
*o1* mutants exhibit abnormal subsidiary cell division. **(A)**. Abnormal subsidiary cells were counted in B73 wildtype, *o1-2995*, *o1-ref* and their corresponding wildtype siblings. Genotypes that are not significantly different at p=0.05 via two-sided t-test are joined noted by the same letter. Different letters indicate significantly different. Methacrylate impressions on leaf 4 of (**B**) B73 (**C**) *o1-ref/+* and (**D**) *o1-ref*, (**E**) *o1-2995/+* and (**F**) *o1-2995*. Abnormal cells are highlighted in brown.

To determine when the asymmetric division of the subsidiary mother cell becomes misoriented, *o1-2995* and *o1-tan62* were crossed to a live cell marker for microtubules (YFP-TUBULIN or CFP-TUBULIN <u>(Mohanty et al. 2009)</u>). Then, three wild-type siblings and three mutants were dissected to reveal the asymmetric division zone and imaged. In wild-type siblings, premitotic microtubule structures were oriented correctly (**Figure 6A**). Similarly, spindles and preprophase bands in *o1-2995* and *o1-tan62* were similar to wild type. (**Figure 6B and 6C**). In *o1-2995* and *o1-tan62* mutants, phragmoplasts were often misoriented (∼60% for both). These data indicate that *o1-2995* and *o1-tan62* exhibit phragmoplast guidance defects, similar to, but more severe in frequency than, *o1* null mutants previously described (Nan et al., 2023).

**Figure 6.**
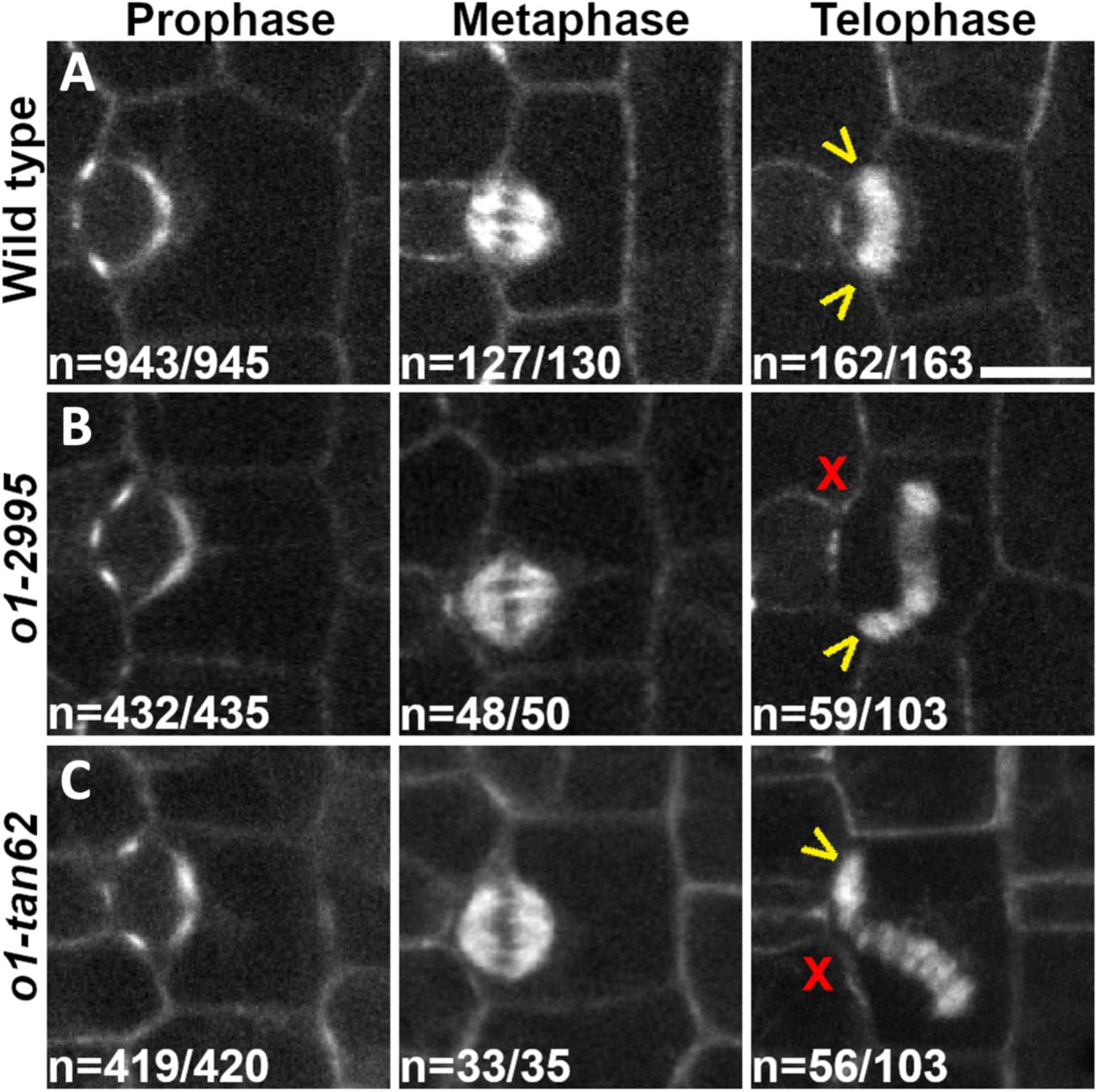
*o1-2995* and *o1-tan62* have phragmoplast guidance defects. Representative micrographs of each cell cycle stage in plants expressing a TUBULIN fluorescent marker. Three plants of *o1-2995, o1-tan62,* and their respective wild-type siblings were analyzed. For prophase, counts indicate total number of cells with normal preprophase bands. Wild type sibling of *o1-2995*: n=442/442 cells; wild-type sibling of *o1-tan62*: n=501/503 cells. For metaphase, counts indicate the total number of cells with normal spindle morphology. Wild type sibling of *o1-2995*: n=80/83 cells; wild-type sibling of *o1-tan62*: n=47/47 cells. For telophase, count in wild type indicates the total number of oriented phragmoplasts. Wild type sibling of *o1-2995*: n=90/91 cells; wild-type sibling of *o1-tan62*: n=72/72 cells. Telophase cell count in *o1-2995* and *o1-tan62* indicates total number of cells with misoriented phragmoplasts. Yellow arrows = oriented phragmoplast. Red X = misoriented phragmoplast. Scale bar = 10 µm.

## Discussion

Similar to *o1* mutants, triple and quadruple mutants in class XI myosins in Arabidopsis generate smaller plants, with reduced cell expansion, short roots, aberrant organelle movement and division plane positioning defects <u>(Peremyslov et al. 2010; Madison et al. 2015)(Abu-Abied et</u> <u>al. 2018; Huang et al. 2024)</u>. Often, myosin XI mutants have shortened root hairs or pollen tubes indicating cell elongation defects <u>(Madison et al. 2015; Peremyslov et al. 2015)</u>. In *Physcomitrium patens,* RNAi directed at the only two myosin XIs generated very small plants with round cells also indicating a critical role in cell elongation <u>(Vidali et al. 2010)</u>. In contrast, single myosin XI mutants (*myosin XI-I*) in Arabidopsis have more subtle phenotypes: nuclear migration defects that generate aberrant division planes during stomatal development that reduce stomatal density <u>(Muroyama et al. 2020)</u> and nuclear shape and movement defects <u>(Tamura et al.</u> <u>2013)</u>.

Defects in nuclear migration/polarization during subsidiary cell development were not detected in the *o1* mutant in maize, but both ER organization and subsidiary cell division positioning were aberrant <u>(Nan et al. 2023; Wang et al. 2012)</u>. The *o1* mutant was originally identified due to the opaque kernel phenotype, which is caused by aberrant protein body accumulation in the endosperm <u>(Wang et al. 2012)</u>. The O1 protein associates with the ER, and also as puncta in the cytoplasm and in the phragmoplast midline <u>(Nan et al. 2023; Wang et al. 2012)</u>. Overexpression of the tail domain disrupts ER motility in tobacco cells (Wang et al., 2012).

Plant class XI myosins are processive motors that dimerize and transport cargo and promote cytoplasmic streaming <u>(Tominaga and Nakano 2012; Tominaga et al. 2003)</u>. Overexpression or ectopic expression of truncated myosins lacking the motor domain has a dominant-negative effect that impairs myosin activity in vivo, potentially due to dimerization <u>(Stephan et al. 2021;</u> <u>Peremyslov et al. 2008; Sparkes et al. 2008; Avisar et al. 2008; Wang et al. 2012; Avisar et al.</u> <u>2009)</u>. As a motor, Arabidopsis Myosin XI-I has unique features, including a high affinity for actin with corresponding low velocity and ATPase activity compared to other myosin XIs <u>(Haraguchi et al. 2016)</u>. However, individual myosins are likely to have different actin binding affinities, cargo, and velocities, and therefore different (but potentially overlapping) cellular functions. These diverse cellular functions may translate into different phenotypes on the tissue, organ and plant level. Previous immunoprecipitation experiments show that O1 interacts with other myosins, including Myosin XI-K, XI-F and XI-G family members as well as multiple myosin VIIIs <u>(Nan et al. 2023)</u>. Notably, *O1* has now been identified in four genetic screens, with three distinct phenotypes <u>(Nan et al. 2023; Wang et al. 2012)</u>. To the best of our knowledge, no other maize gene encoding a myosin has been identified in a genetic screen. This implies that O1’s function may be unique - either because of its intrinsic properties, such as binding affinity or cargo - or because it heterodimerizes with many different myosins. O1 also interacts with actin binding proteins, heat shock interacting proteins and kinesins that are related to PHRAGMOPLAST ORIENTING KINESIN1 (POK1) and POK2 <u>(Nan et al. 2023; Wang et al. 2012; Müller et al. 2006)</u>. Determining the scope of dimerization and cargos across different myosins in future studies may help elucidate O1 and myosin function in general.

Our current hypothesis is that both missense mutant alleles described in this paper generate proteins with impaired function of a motor domain activity, such as actin binding or the ATPase activity required for movement. We predict that O1-2995 and O1-TAN62 mutant proteins, despite predicted defects in motor domain function, would still be capable of interacting with multiple other proteins, including both myosin XIs and myosin VIIIs <u>(Nan et al. 2023)</u>, and cargoes, as outlined in our speculative model (**Figure 7**). The *o1-2995* mutant is recessive: no antimorph effect is seen in either simple (*O1/o1-2995*) or compound (*o1-ref/o1-2995*) heterozygous plants. We speculate that the recessive nature of *o1-2995* reflects insufficient expression levels of the mutant form of the protein. Thus, the more severe mutant phenotype of homozygous *o1-2995* or *o1-tan-62* relative to *o1* null mutants could be due to O1 mutant proteins binding to and reducing activity of other myosins (**Figure 7**) or even other binding partners, potentially including cargo. It is plausible that the antimorph effects are only observed in homozygous *o1-2295* mutants due to dosage effects, i.e., increased expression of *o1-2995* coupled with the absence of functional O1 may sufficiently increase the relative abundance of poisoned O1-containing protein complexes to enhance the plant growth and cell division positioning phenotypes.

**Figure 7.**
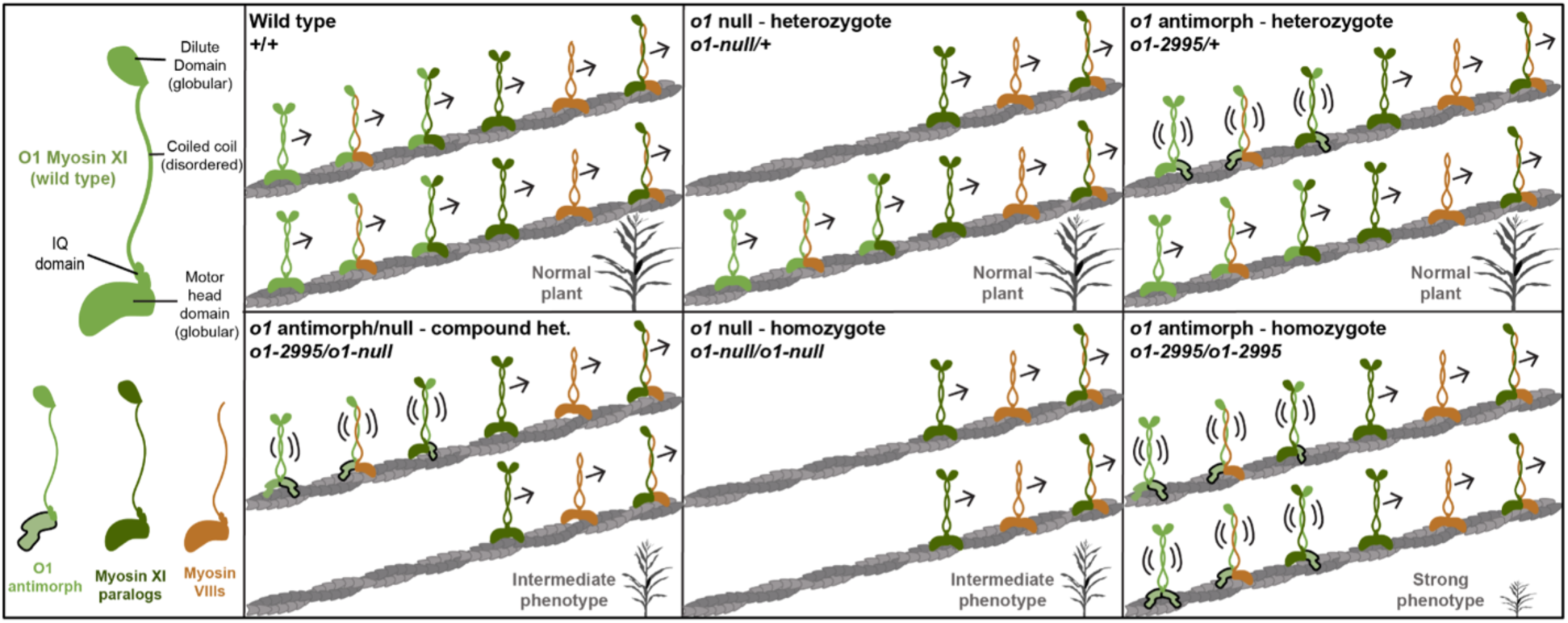
Model of O1 cellular function across different genotypes. Myosin VIII (brown) and XI (green) polypeptide monomers contain a motor head, a neck region containing IQ domains and a tail region with coiled coil domains involved in cargo binding. Myosin XI monomers (including O1, light green) also contain a dilute domain in the tail region. The defective motor head of the O1 antimorph monomer is indicated by a misshapen motor with a solid outline. Each main panel illustrates one genotype and its expressed myosin dimers bound to actin microfilaments (gray), and whether the dimers are predicted to have functional motor activity (arrows) or not (double parentheses). Though depicted as such, antimorph-containing dimers may or may not bind to actin. **Top Left** In wild type plants, O1 homodimerizes and heterodimerizes with other Myosin XI isoforms and with Myosin VIII. Relative numbers of hetero- and homodimers *in vivo* is unknown; this panel illustrates two “copies” of each dimer type. Each myosin dimer will have associated cargo (not shown), which could be specific or shared with other dimers. **Top Center** In *o1* null heterozygotes, the frequency of dimers containing O1 decreases, but all dimer types are present. The plants are phenotypically wild type. **Bottom Center** Complete loss of O1 in *o1* null mutants eliminates any complex containing O1, leading to fewer dimer types and an intermediate plant phenotype. **Top Right** In *o1* antimorph heterozygotes, O1-2995 and O1-tan62 proteins have defective head domains, leading to “poisoned” complexes. Nonetheless, these alleles behave as fully recessive and not as dominant negatives, i.e. heterozygotes are phenotypically normal, plausibly because some non-poisoned homo- and hetero-dimers form which are sufficient for function. **Bottom Right** In antimorph homozygotes, there are only non-functional O1 homodimers and more poisoned heterodimers, leading to a stronger phenotype than the null mutants. This may be due in part to disabling of functional complexes (i.e., binding to other myosins) or to sequestering cargo (i.e., binding to other proteins, not shown). **Bottom Left** Plants that have one antimorph allele and one null allele have an intermediate phenotype similar to the null homozygote, suggesting a dosage requirement for poisoned complexes to manifest a strong phenotype.

Severity of the subsidiary cell division defects correlated with defects in plant height, but the relationship between cell division defects and plant height is unclear. One potential hypothesis is that gas exchange defects might lead to short plants. However, while aberrant subsidiary cells disrupt the closure of guard cells, either mild or no gas exchange defects were observed in other mutants with aberrant subsidiary cells including *pangloss1 (pan1), pan2,* or the *pan1 pan2* double mutant <u>(Liu et al. 2024)</u>. Alternative and more plausible hypotheses posit that O1 has multiple functions: one required for cell expansion and another that affects subsidiary cell division. Cell expansion defects could be enhanced by additional mechanical stresses generated via aberrantly shaped cells caused by division positioning defects <u>(Sampathkumar et al. 2014)</u>.

## Materials and methods

### Genetic stocks and phenotypic characterization

The *rsl*-12.2995* (*o1-2995*) mutant allele originated from a *ramosa1* (*ra1*) modifier screening population generated via ethyl methane sulfonate (EMS) pollen mutagenesis. *ra1-63* was backcrossed six times to Mo17 to create a homozygous mutant stock. Pollen from mutants was collected, treated with 0.06% EMS in paraffin oil as described <u>(Weeks 2013)</u> and used to pollinate the same genotype. We screened for dominant modifiers in the M1 generation, and then the remaining plants were self-fertilized. The resulting M2 populations were screened for recessive modifiers, where we identified *rsl*-12.2995* (*o1-2995*) as a mutant that segregated 3:1 and displayed short plant stature with suppressed inflorescence branching.

To understand its penetrance and expressivity in diverse genetic backgrounds we backcrossed the *o1-2995* allele to B73, Mo17, and W22 inbred lines at least five times. B73-introgressed stocks for other mutants were sourced from the authors’ labs for *ra1-R* (EV), *ra2-R* (EV), *o1-ref* (MF) and *o1-N1242A* (MF), or from David Jackson (Cold Spring Harbor Laboratory) for *ra3-ref*.

Alleles of *o1-ref* and *o1-N1242A* were obtained from the Maize Genetics Cooperation Stock Center, and introgressed into B73 four times before crossing to *o1-2995*.

*o1-tangled62* (*o1-tan62*) was identified from an EMS mutagenesis performed on B73 kernels. The M0 generation was open-pollinated, followed by self-fertilization of the M1 generation. Glue impressions of leaf 2 or 3 in the M2 generation were examined for division plane positioning defects. *o1-tan62* was backcrossed to B73 at least three times before sequencing the *O1* locus.

### Map-based cloning, sequencing, linkage analysis, and complementation test

To identify the causative mutation lesion responsible for the *rsl*-12.2995* (*o1-2995*) mutant phenotype (Zebosi 2022), we used map-based cloning methods with an F2 mapping population developed between Mo17 and B73.

A bulked segregant analysis using genotyping-by-sequencing (BSA-GBS) method was devised and implemented as follows. A [B73 x Mo17] F2 mapping population was sown and leaf tissue samples collected as two pools bulked from mutant (n=12) and normal sibling (n=26) plants. 150-300 leaf punches (6 mm) were collected for each pool, with an equal number of punches collected from each individual plant in the bulked sample. High molecular weight genomic DNA was extracted using a urea-based protocol <u>(Chen and Dellaporta 1994)</u>. Genotyping-by-sequencing <u>(Elshire et al. 2011)</u> was then adapted and applied to the DNA extracted from the two bulked pools (see <u>(Kokulapalan 2018)</u> for more detail). Briefly, genomic DNAs were quantified using the Promega Quantifluor dsDNA system, and used in the BSA-GBS bench protocol consisting of three main steps. Namely, restriction digestion using ApeKI, ligation of a barcoded and a common adapter, and PCR amplification and cleanup to construct the barcoded sequencing library. Sequencing was performed using 100-bp single-end sequencing with Illumina HiSeq 2500 (Iowa State University, DNA facility). Sequencing Reads were de-multiplexed, separated into independent files for each bulked pool and clipped, all using fastq-mcf <u>(Aronesty 2013)</u>. Reads were aligned to maize B73 RefGen_AGPv4 using minimap2 <u>(Jiao</u> <u>et al. 2017; Li 2018)</u>. Euclidean distance (ED) between the normal and mutant pools was calculated using nucleotide composition and used to calculate ED4 to suppress noise. ED4 distances were plotted on a Manhattan plot using ggplot2 to visualize differences between the pools <u>(Wickham 2016)</u>. (BSA-GBS) roughly mapped the *o1-2995* mutation to the long arm of chromosome 4 (**Supplementary** Figure 2), within a peak containing approximately 989 genes.

Using publicly available molecular markers, we further mapped the mutation to a region between markers IDP2 (18 recombinants, 2.24 cM) and IDP8018 (41 recombinants, 5.11 cM), and then with public SNPs to between BZ2995_2 (8 recombinants, 1 cM) and BZ2995_1 (2 recombinants, 0.25 cM), comprising a physical distance of 1.5Mb and 28 genes (**Figure 2A**). To further narrow down the location of the causal mutation, we used two fine-mapping populations (F2 and F1BC1) constructed by crossing *o1-2995* mutants in the original Mo17 genetic background to the B73 inbred line. DNA was extracted as previously described <u>(Zebosi et al. 2025)</u>. Using publicly available insertion/deletion polymorphisms between B73 and Mo17, we designed new markers to reduce the fine-mapping to the interval between IDP2 (18/802, 2.24 cM) and IDP8018 (41/802, 5.11 cM). To further narrow the interval, single nucleotide polymorphisms (SNPs) were used to design new markers that localized the mutation to a 1.5Mb interval between BZ2995_6 (8/802, 1 cM) and BZ2995_1 (2/802, 0.25 cM) with 28 genes (**Figure 2A**). In non-repetitive regions between markers BZ2995_6 and BZ2995_1, B73 and Mo17 contained no DNA sequence polymorphisms to advance the fine-mapping process. We therefore performed whole-genome sequencing (WGS) of a single mutant plant to identify EMS-induced SNPs in the interval for marker development and identifying potential candidate genes. The library was prepared and sequenced on an Illumina HiSeq by Novogene Inc., UC Davis, California, to generate 50Gb of 150-bp paired-end sequencing data. The raw reads were subjected to quality control using FASTQC <u>(Andrews 2010)</u> and adapters, and reads with low base quality were trimmed using Trimmomatic version 0.32 <u>(Bolger et al. 2014)</u>. The trimmed reads were aligned to the Mo17 genome using the ‘mem’ algorithm in BWA-MEM version 0.7.17 <u>(Li and Durbin</u> <u>2010)</u>, and the resulting sequence alignment SAM files were converted to BAM files and filtered for uniquely mapped reads using Samtools <u>(Li et al. 2009)</u>. SNP calling against the Mo17 genome was performed using the Samtools mpileup command, and homozygous non-Mo17 or EMS SNPs (G>A and C > T) in the 1.5Mb interval were identified. We identified 3 genes with exonic EMS SNPs, GRMZM2G521481, GRMZM2G091143, and GRMZM2G449909 (*Opaque1/O1*).

From the EMS-like SNPs we developed new markers and used them in F2 populations consisting of 2289 plants (both wildtype and mutants) to find six recombination events between the three candidate genes. Left-side recombinants (3 and 17) and right-side recombinants (9 and 16) suggested that the causal mutation was within a physical distance of ∼ 0.6Mb that contained only one gene, the previously cloned *Opaque1* (*O1*) (**Figure 2B**). We therefore phenotyped kernels from our segregating populations, and found an opaque kernel phenotype co-segregated with the *o1-2995* mutants. We verified the six recombinants by test crossing each with *o1-2995* mutants and phenotyping the progeny for kernel opaqueness and plant height. To validate *Opaque1* as the causal locus, we performed a complementation test by crossing *o1-2995* mutant to two *o1* mutant alleles (*o1-ref/+* and *o1-N1243/+*).

To sequence the *O1* coding sequence in *tan62* mutants, RNA was extracted from young leaf tissue of *o1-tan62* homozygous mutants and wild-type plants using the RNeasy Plant Mini Kit (QIAGEN) and reverse transcribed into cDNA using the QuantiTect Reverse Transcription Kit (QIAGEN). Then, the cDNA was amplified into two halves and sequenced using primers in Supplementary Table 1. PCR products were run on a gel and bands were gel purified using the QIAquick Gel Extraction Kit (QIAGEN). Seven PCR products spanning the entire coding sequence were sent for sanger sequencing at the Genomics Core Facility (University of California, Riverside). *o1-tan62* and wild-type sequences were compared to each other and GRMZM2G449909 (*O1*) by using A plasmid Editor <u>(Davis and Jorgensen 2022)</u> and NCBI BLAST <u>(Altschul et al. 1990)</u>. DNA from an additional 3 *tan62* mutants was extracted, and sequenced PCR products confirmed the mutation. Complementation test crosses between *o1-tan62* and *o1-N1242A* homozygous mutants were done in the field at the University of California, Riverside in the summer of 2021. Approximately 50 kernels each from 11 independent crosses were assessed for and had opaque kernel phenotype confirming that *tan62* was allelic to *o1*.

### Plant growth conditions

Field-grown plants were grown in the summer maize nursery at Curtiss Farms, Iowa State University in Ames, Iowa or at University of California, Riverside (UCR) Agricultural Operations. UCR greenhouse growing conditions: maize kernels were planted in 3.8 liter pots with soil (20% peat, 50% bark, 10% perlite, and 20% medium vermiculite) supplemented with magnesium nitrate (50 ppm N and 45 ppm Mg), calcium nitrate (75 ppm N and 90 ppm Ca), and Osmocote Classic 3-4 M (NPK 14-14-14%, AICL SKU #E90550) with standard greenhouse conditions (31-33°C daytime temperature with supplemental lighting from 17:00-21:00 PM at ∼ 400 µEm^-2^s^-1^) <u>(Uyehara et al. 2024)</u>

### Phylogenetic and gene expression analysis

Myosin amino acid sequences of Arabidopsis, maize, rice, Setaria, sorghum, and Physcomitrella were obtained from Phytozome <u>(Goodstein et al. 2012)</u>. Sequence alignments were performed using ClustalW2 <u>(Larkin et al. 2007)</u> and Mesquite software (Maddison and Maddison, 2011). Approximate maximum-likelihood phylogenetic trees were constructed as described by <u>(Zebosi</u> <u>et al. 2024)</u>. Raw RNA sequence reads from several tissues were downloaded from the short read archive (https://www.ncbi.nlm.nih.gov/sra) and reads were analyzed as described (Zebosi et al., 2024).

### Protein analysis

Membrane proteins were extracted from the basal 0.5-3 cm of leaves from the 4-leaf stage and separated by SDS-PAGE, according to <u>(Facette et al. 2015)</u>. The previously described rabbit antibody was generated by injection of O1-specific peptides <u>(Nan et al. 2023)</u> and used at 0.66 micrograms/ml.

### Phenotypic characterization

Phenotypic characterization was done using the *o1-2995* allele or the *o1-tan62* allele introgressed in B73 compared to their corresponding wild-type siblings. Agronomic traits such as plant height, total leaf number, leaf length and width, tassel length, tassel, and ear branch number were characterized using segregating populations. Plant height measurements were taken by measuring from the soil surface to the tip of the tassel at maturity. Total leaf number was tracked from germination and measured in mature plants. Peduncle length was measured from the flag leaf node to the lowest tassel branch. Tassel length was measured from the lowest tassel branch to the tip of the tassel. Internode length and diameter, and peduncle lengths were measured with a tape measure. For internode length, measurements were taken starting from the first internode closest to the tassel proceeding basipetally to the 12th internode from the tassel.

### Phloroglucinol-HCl staining

For Phloroglucinol-HCl staining, midribs from adult leaf 10 were hand-sectioned with a double-edge razor, and sections were stained using a Phloroglucinol-HCl staining solution <u>(Strable et al.</u> <u>2017)</u>. The stained sections were observed under a dissecting microscope (Leica MZ125) and imaged using a digital camera.

### Leaf epidermal impressions

Leaf epidermal impressions of the abaxial surface were produced from mid-length between the ligule and leaf tip and mid-way between the leaf margin and the mid-vein using cyanoacrylate glue as described <u>(Allsman et al. 2019)</u>.. The slides with the epidermal imprints were observed under the Olympus BX60 light microscope and imaged using a digital camera (Jenoptik C5). The size of epidermal cells and the number of aberrant subsidiary cells were quantified from three random viewable fields using ImageJ <u>(Rueden et al. 2017)</u>. For *o1-tan62* mutants and wild-type siblings, leaf epidermal impressions were taken from the abaxial side of the second leaf. Epidermal imprints were imaged with a light compound microscope (Nikon) with digital microscope camera attachment (AmScope MD130).

### Confocal Microscopy

Images were taken of the asymmetric divisions of the subsidiary mother cell using a Yokogawa W1 spinning disk on a Nikon Eclipse TE inverted stand microscope with an EM-CCD camera (Hamamatsu 9100c). The ASI Peizo stage and 3 axis DC servo motor controllers were controlled using Micromanager software (www.micromanager.org). Solid-state Obis lasers (power from 40 to 100 mW) were used in combination with standard emission filters (Chroma Technology). For CFP-ꞵ-TUBULIN, a 445 laser with emission filter of 480/40 was used. For YFP-*ɑ*-TUBULIN, a 514 laser with emission filter 540/30 was used. *o1-2995*, *o1-tan62*, and their wild-type siblings, expressing CFP-ꞵ-TUBULIN or YFP-*ɑ*-TUBULIN <u>(Mohanty et al. 2009)</u>, were grown in the greenhouse under standard conditions as described (Uyehara et al. 2024). Four week-old plants were dissected to an emerging leaf with ligule height < 2 mm, and the asymmetric dividing zone (within 1.5 cm from the ligule) was mounted in a rose chamber <u>(Rasmussen 2016)</u>. Micrographs of asymmetric divisions of the subsidiary mother cell were captured along the entirety of the section and subsequently analyzed using the FIJI version of ImageJ <u>(Schindelin et al. 2012)</u>.

## Supporting information

All Supplemental Figures

## Data Analysis and Figure Preparation

Graphs and statistics were done using the R software environment for statistical computing and graphics (https://www.R-project.org/) and RStudio software https://posit.co/ using several packages. Some figures were assembled using the Gnu Image Manipulation Program (Gimp, version 2.10.38, https://www.gimp.org).

## Accession numbers

*Opaque1 (O1):* GRMZM2G449909 (B73 RefGen_v3), Zm00001d052110 (Zm-B73-REFERENCE-GRAMENE-4.0), Zm00001eb193160 (Zm-B73-REFERENCE-NAM-5.0)

*Ramosa1 (Ra1):* GRMZM2G361210 (B73 RefGen_v3), Zm00001d034642 (Zm-B73-REFERENCE-GRAMENE-4.0), Zm00001eb062570 (Zm-B73-REFERENCE-NAM-5.0)

*Ramosa2 (Ra2):* AC233943.1_FG002 (B73 RefGen_v3), Zm00001d039694 (Zm-B73-REFERENCE-GRAMENE-4.0), Zm00001eb123060 (Zm-B73-REFERENCE-NAM-5.0)

*Ramosa3 (Ra3):* GRMZM2G014729 (B73 RefGen_v3), Zm00001d022193 (Zm-B73-REFERENCE-GRAMENE-4.0), Zm00001eb327910 (Zm-B73-REFERENCE-NAM-5.0)

## Acknowledgments

We want to thank Connor Hamers (ISU), Jack Schwickerath (ISU), Nicole Essner (ISU), David Wetovic (UCR), and Lindy Allsman (UCR) for their help with the summer fieldwork, and the Curtiss Farms (ISU) and Agricultural Operations (UCR) staff for maize genetics nursery efforts. We thank Colin Finnegan (ISU) and Fred Roger Namanda (ISU) for help with plant phenotyping, Erica Unger-Wallace (ISU) for training in laboratory methods and Professor David Nelson (UCR) for discussing genetics. Thanks to the Maize Genetics Cooperation Stock Center for providing maize kernels. This work was supported by the NSF-PGRP 1238202 to E.V., NSF-CAREER 1942734 and NSF-2426623 to C.G.R., NSF NRT Plants3D (DBI-1922642) to S.E.M. and G.S.S.

## Author contributions

E.V. conceptualized the project; B.Z., S.E.M, and E.V. curated data; B.Z. and S.E.M. performed formal analysis; S.E.M., G.S.B., M.F., C.R. and E.V. acquired funding; B.Z., S.E.M., K.W., J.S., G.S.B., N.B.B., M.F. and E.V. performed experiments and/or collected data; C.R. and E.V. administered and supervised the project; M.F., C.R. and E.V. provided material resources for study; K.W. developed software; B.Z., S.E.M. and K.W. performed validation; B.Z., S.E.M., K.W., N.B.B., M.F., C.R. and E.V. prepared figures for visualization; B.Z. wrote the original draft of the manuscript; B.Z., S.E.M., K.W., M.F., C.R. and E.V. reviewed and edited the manuscript.

## Literature Cited

Abu-Abied M, Belausov E, Hagay S, Peremyslov V, Dolja V, Sadot E (2018) Myosin XI-K is involved in root organogenesis, polar auxin transport, and cell division. J Exp Bot 69: 2869–2881

Allsman LA, Dieffenbacher RN, Rasmussen CG (2019) Glue impressions of maize leaves and their use in classifying mutants. Bio-protocol 9: e3209

Altschul SF, Gish W, Miller W, Myers EW, Lipman DJ (1990) Basic local alignment search tool. J Mol Biol 215: 403–410

Andrews S (2010) FastQC: quality control tool high throughput sequence data. Babraham Bioinformatics

Aronesty E (2013) Comparison of sequencing utility programs. Open Bioinforma J 7: 1–8

Avisar D, Abu-Abied M, Belausov E, Sadot E, Hawes C, Sparkes IA (2009) A comparative study of the involvement of 17 Arabidopsis myosin family members on the motility of Golgi and other organelles. Plant Physiol 150: 700–709

Avisar D, Prokhnevsky AI, Makarova KS, Koonin EV, Dolja VV (2008) Myosin XI-K Is Required for Rapid Trafficking of Golgi Stacks, Peroxisomes, and Mitochondria in Leaf Cells of Nicotiana benthamiana. Plant Physiol 146: 1098–1108

Barazesh S, McSteen P (2008) Barren inflorescence1 functions in organogenesis during vegetative and inflorescence development in maize. Genetics 179: 389–401

Bolger AM, Lohse M, Usadel B (2014) Trimmomatic: a flexible trimmer for Illumina sequence data. Bioinformatics 30: 2114–2120

Bortiri E, Chuck G, Vollbrecht E, Rocheford T, Martienssen R, Hake S (2006) ramosa2 encodes a LATERAL ORGAN BOUNDARY domain protein that determines the fate of stem cells in branch meristems of maize. Plant Cell 18: 574–585

Chen J, Dellaporta S (1994) Urea-based Plant DNA Miniprep. The Maize Handbook. Springer New York, New York, NY, pp 526–527

Chocano-Coralla EJ, Vidali L (2024) Myosin XI, a model of its conserved role in plant cell tip growth. Biochem Soc Trans 52: 505–515

Claeys H, Vi SL, Xu X, Satoh-Nagasawa N, Eveland AL, Goldshmidt A, Feil R, Beggs GA, Sakai H, Brennan RG, et al (2019) Control of meristem determinacy by trehalose 6-phosphate phosphatases is uncoupled from enzymatic activity. Nat Plants 5: 352–357

Davis MW, Jorgensen EM (2022) ApE, A Plasmid Editor: A Freely Available DNA Manipulation and Visualization Program. Frontiers in Bioinformatics. doi: 10.3389/fbinf.2022.818619

Elshire RJ, Glaubitz JC, Sun Q, Poland JA, Kawamoto K, Buckler ES, Mitchell SE (2011) A robust, simple genotyping-by-sequencing (GBS) approach for high diversity species. PLoS One 6: e19379

Eveland AL, Goldshmidt A, Pautler M, Morohashi K, Liseron-Monfils C, Lewis MW, Kumari S, Hiraga S, Yang F, Unger-Wallace E, et al (2014) Regulatory modules controlling maize inflorescence architecture. Genome Res 24: 431–443

Facette MR, Park Y, Sutimantanapi D, Luo A, Cartwright HN, Yang B, Bennett EJ, Sylvester AW, Smith LG (2015) The SCAR/WAVE complex polarizes PAN receptors and promotes division asymmetry in maize. Nat Plants 1: 14024

Foth BJ, Goedecke MC, Soldati D (2006) New insights into myosin evolution and classification. Proc Natl Acad Sci U S A 103: 3681–3686

Gallavotti A, Long JA, Stanfield S, Yang X, Jackson D, Vollbrecht E, Schmidt RJ (2010) The control of axillary meristem fate in the maize ramosa pathway. Development 137: 2849–2856

Gallavotti A, Zhao Q, Kyozuka J, Meeley RB, Ritter MK, Doebley JF, Pè ME, Schmidt RJ (2004) The role of barren stalk1 in the architecture of maize. Nature 432: 630– 635

Galli M, Liu Q, Moss BL, Malcomber S, Li W, Gaines C, Federici S, Roshkovan J, Meeley R, Nemhauser JL, et al (2015) Auxin signaling modules regulate maize inflorescence architecture. Proc Natl Acad Sci U S A 112: 13372–13377

Golomb L, Abu-Abied M, Belausov E, Sadot E (2008) Different subcellular localizations and functions of Arabidopsis myosin VIII. BMC Plant Biol 8: 3

Goodstein DM, Shu S, Howson R, Neupane R, Hayes RD, Fazo J, Mitros T, Dirks W, Hellsten U, Putnam N, et al (2012) Phytozome: a comparative platform for green plant genomics. Nucleic Acids Res 40: D1178–86

Haraguchi T, Tominaga M, Nakano A, Yamamoto K, Ito K (2016) Myosin XI-I is Mechanically and Enzymatically Unique Among Class-XI Myosins in Arabidopsis. Plant Cell Physiol 57: 1732–1743

Huang CH, Peng FL, Lee Y-RJ, Liu B (2024) The microtubular preprophase band recruits Myosin XI to the cortical division site to guide phragmoplast expansion during plant cytokinesis. Dev Cell. doi: 10.1016/j.devcel.2024.05.015

Jiao Y, Peluso P, Shi J, Liang T, Stitzer MC, Wang B, Campbell MS, Stein JC, Wei X, Chin C-S, et al (2017) Improved maize reference genome with single-molecule technologies. Nature 546: 524–527

Kokulapalan W (2018) Experimental and computational methods to assign gene function to maize genes.

Lambert RJ, Johnson RR (1978) Leaf angle, tassel morphology, and the performance of maize hybrids^1^. Crop Sci 18: 499–502

Larkin MA, Blackshields G, Brown NP, Chenna R, McGettigan PA, McWilliam H, Valentin F, Wallace IM, Wilm A, Lopez R, et al (2007) Clustal W and Clustal X version 2.0. Bioinformatics 23: 2947–2948

Li H (2018) Minimap2: pairwise alignment for nucleotide sequences. Bioinformatics 34: 3094–3100

Li H, Durbin R (2010) Fast and accurate long-read alignment with Burrows-Wheeler transform. Bioinformatics 26: 589–595

Li H, Handsaker B, Wysoker A, Fennell T, Ruan J, Homer N, Marth G, Abecasis G, Durbin R, 1000 Genome Project Data Processing Subgroup (2009) The Sequence Alignment/Map format and SAMtools. Bioinformatics 25: 2078–2079

Liu L, Ashraf MA, Morrow T, Facette M (2024) Stomatal closure in maize is mediated by subsidiary cells and the PAN2 receptor. New Phytol 241: 1130–1143

Madison SL, Buchanan ML, Glass JD, McClain TF, Park E, Nebenführ A (2015) Class XI Myosins Move Specific Organelles in Pollen Tubes and Are Required for Normal Fertility and Pollen Tube Growth in Arabidopsis. Plant Physiol 169: 1946–1960

Maddison and Maddison (2011) Mesquite: a modular system for evolutionary analysis. Evolution, 1103–1118. Retrieved from http://mesquiteproject.org

Madison SL, Nebenführ A (2013) Understanding myosin functions in plants: are we there yet? Curr Opin Plant Biol 16: 710–717

McSteen P, Hake S (2001) Barren inflorescence2 regulates axillary meristem development in the maize inflorescence. Development 128: 2881–2891

McSteen P, Malcomber S, Skirpan A, Lunde C, Wu X, Kellogg E, Hake S (2007) barren inflorescence2 Encodes a co-ortholog of the PINOID serine/threonine kinase and is required for organogenesis during inflorescence and vegetative development in maize. Plant Physiol 144: 1000–1011

Mohanty A, Luo A, DeBlasio S, Ling X, Yang Y, Tuthill DE, Williams KE, Hill D, Zadrozny T, Chan A, et al (2009) Advancing cell biology and functional genomics in maize using fluorescent protein-tagged lines. Plant Physiol 149: 601–605

Mühlhausen S, Kollmar M (2013) Whole genome duplication events in plant evolution reconstructed and predicted using myosin motor proteins. BMC Evol Biol 13: 202

Müller S, Han S, Smith LG (2006) Two kinesins are involved in the spatial control of cytokinesis in Arabidopsis thaliana. Curr Biol 16: 888–894

Muroyama A, Gong Y, Bergmann DC (2020) Opposing, Polarity-Driven Nuclear Migrations Underpin Asymmetric Divisions to Pattern Arabidopsis Stomata. Curr Biol. doi: 10.1016/j.cub.2020.08.100

Nan Q, Liang H, Mendoza J, Liu L, Fulzele A, Wright A, Bennett EJ, Rasmussen CG, Facette MR (2023) The OPAQUE1/DISCORDIA2 myosin XI is required for phragmoplast guidance during asymmetric cell division in maize. Plant Cell 35: 2678–2693

Nebenführ A, Dixit R (2018) Kinesins and Myosins: Molecular Motors that Coordinate Cellular Functions in Plants. Annu Rev Plant Biol 69: 329–361

Neuffer MG, Jones L, Zuber MS (1968) The mutants of maize. doi: 10.2135/1968.mutantsofmaize

Olatunji D, Clark NM, Kelley DR (2023) The class VIII myosin ATM1 is required for root apical meristem function. Development 2022.11.30.518567

Pendleton JW, Smith GE, Winter SR, Johnston TJ (1968) Field investigations relationships leaf angle corn (Zea mays L.) grain yield apparent photosynthesis. Agronomy Journal 60: 422–424

Peremyslov VV, Cole RA, Fowler JE, Dolja VV (2015) Myosin-Powered Membrane Compartment Drives Cytoplasmic Streaming, Cell Expansion and Plant Development. PLoS One 10: e0139331

Peremyslov VV, Mockler TC, Filichkin SA, Fox SE, Jaiswal P, Makarova KS, Koonin EV, Dolja VV (2011) Expression, splicing, and evolution of the myosin gene family in plants. Plant Physiol 155: 1191–1204

Peremyslov VV, Prokhnevsky AI, Avisar D, Dolja VV (2008) Two class XI myosins function in organelle trafficking and root hair development in Arabidopsis. Plant Physiol 146: 1109–1116

Peremyslov VV, Prokhnevsky AI, Dolja VV (2010) Class XI myosins are required for development, cell expansion, and F-Actin organization in Arabidopsis. Plant Cell 22: 1883– 1897

Rasmussen CG (2016) Using Live-Cell Markers in Maize to Analyze Cell Division Orientation and Timing. *In* M-C Caillaud, ed, Plant Cell Division: Methods and Protocols. Springer New York, New York, NY, pp 209–225

Reddy AS, Day IS (2001) Analysis of the myosins encoded in the recently completed Arabidopsis thaliana genome sequence. Genome Biol 2: RESEARCH0024

Richards TA, Cavalier-Smith T (2005) Myosin domain evolution and the primary divergence of eukaryotes. Nature 436: 1113–1118

Rueden CT, Schindelin J, Hiner MC, DeZonia BE, Walter AE, Arena ET, Eliceiri KW (2017) ImageJ2: ImageJ for the next generation of scientific image data. BMC Bioinformatics 18: 529

Sampathkumar A, Krupinski P, Wightman R, Milani P, Berquand A, Boudaoud A, Hamant O, Jönsson H, Meyerowitz EM (2014) Subcellular and supracellular mechanical stress prescribes cytoskeleton behavior in Arabidopsis cotyledon pavement cells. Elife 3: e01967

Satoh-Nagasawa N, Nagasawa N, Malcomber S, Sakai H, Jackson D (2006) A trehalose metabolic enzyme controls inflorescence architecture in maize. Nature 441: 227–230

Schindelin J, Arganda-Carreras I, Frise E, Kaynig V, Longair M, Pietzsch T, Preibisch S, Rueden C, Saalfeld S, Schmid B, et al (2012) Fiji: an open-source platform for biological-image analysis. Nat Methods 9: 676–682

Sparkes IA, Teanby NA, Hawes C (2008) Truncated myosin XI tail fusions inhibit peroxisome, Golgi, and mitochondrial movement in tobacco leaf epidermal cells: a genetic tool for the next generation. J Exp Bot 59: 2499–2512

Stephan L, Jakoby M, Das A, Koebke E, Hülskamp M (2021) Unravelling the molecular basis of the dominant negative effect of myosin XI tails on P-bodies. PLoS One 16: e0252327

Strable J, Wallace JG, Unger-Wallace E, Briggs S, Bradbury PJ, Buckler ES, Vollbrecht E (2017) Maize YABBY Genes drooping leaf1 and drooping leaf2 Regulate Plant Architecture. Plant Cell 29: 1622–1641

Tamura K, Iwabuchi K, Fukao Y, Kondo M, Okamoto K, Ueda H, Nishimura M, Hara-Nishimura I (2013) Myosin XI-i links the nuclear membrane to the cytoskeleton to control nuclear movement and shape in Arabidopsis. Curr Biol 23: 1776–1781

Tian J, Wang C, Xia J, Wu L, Xu G, Wu W, Li D, Qin W, Han X, Chen Q, et al (2019) Teosinte ligule allele narrows plant architecture and enhances high-density maize yields. Science 365: 658–664

Tominaga M, Kojima H, Yokota E, Orii H, Nakamori R, Katayama E, Anson M, Shimmen T, Oiwa K (2003) Higher plant myosin XI moves processively on actin with 35 nm steps at high velocity. EMBO J 22: 1263–1272

Tominaga M, Nakano A (2012) Plant-Specific Myosin XI, a Molecular Perspective. Front Plant Sci 3: 211

Ueda H, Tamura K, Hara-Nishimura I (2015) Functions of plant-specific myosin XI: from intracellular motility to plant postures. Curr Opin Plant Biol 28: 30–38

Uyehara AN, Diep BN, Allsman L, Gayer SG, Martinez SE, Kim JJ, Agarwal S, Rasmussen CG (2024) De novo TANGLED1 recruitment from the phragmoplast to aberrant cell plate fusion sites in maize. J Cell Sci. doi: 10.1242/jcs.262097

Vidali L, Burkart GM, Augustine RC, Kerdavid E, Tüzel E, Bezanilla M (2010) Myosin XI is essential for tip growth in Physcomitrella patens. Plant Cell 22: 1868–1882

Vollbrecht E, Springer PS, Goh L, Buckler ES 4th, Martienssen R (2005) Architecture of floral branch systems in maize and related grasses. Nature 436: 1119–1126

Walley JW, Sartor RC, Shen Z, Schmitz RJ, Wu KJ, Urich MA, Nery JR, Smith LG, Schnable JC, Ecker JR, et al (2016) Integration of omic networks in a developmental atlas of maize. Science 353: 814–818

Wang G, Wang F, Wang G, Wang F, Zhang X, Zhong M, Zhang J, Lin D, Tang Y, Xu Z, et al (2012) Opaque1 encodes a myosin XI motor protein that is required for endoplasmic reticulum motility and protein body formation in maize endosperm. Plant Cell 24: 3447–3462

Weeks R (2013) Inflorescence branching in maize: A quantitative genetics approach to identifying key players in the inflorescence development pathway.

Wickham H (2016) Ggplot2, 2nd ed. doi: 10.1007/978-3-319-24277-4

Zebosi B (2022) Functional characterization of ramosa suppressor locus-12.2995 that shows the opaque1 gene regulates plant architecture in maize. search.proquest.com

Zebosi B, Ssengo J, Geadelmann LF, Unger-Wallace E, Vollbrecht E (2025) An effective and safe maize seed chipping protocol using clipping pliers with applications in small-scale genotyping and marker-assisted breeding. Bio Protoc 15: e5200

Zebosi B, Vollbrecht E, Best NB (2024) Brassinosteroid biosynthesis and signaling: Conserved and diversified functions of core genes across multiple plant species. Plant Commun 5: 100982

Zhang W, Cai C, Staiger CJ (2019) Myosins XI Are Involved in Exocytosis of Cellulose Synthase Complexes. Plant Physiol 179: 1537–1555

