## Supplementary material for "Recessive antimorph alleles reveal novel functions of the OPAQUE1 myosin XI in maize": All Supplemental Figures

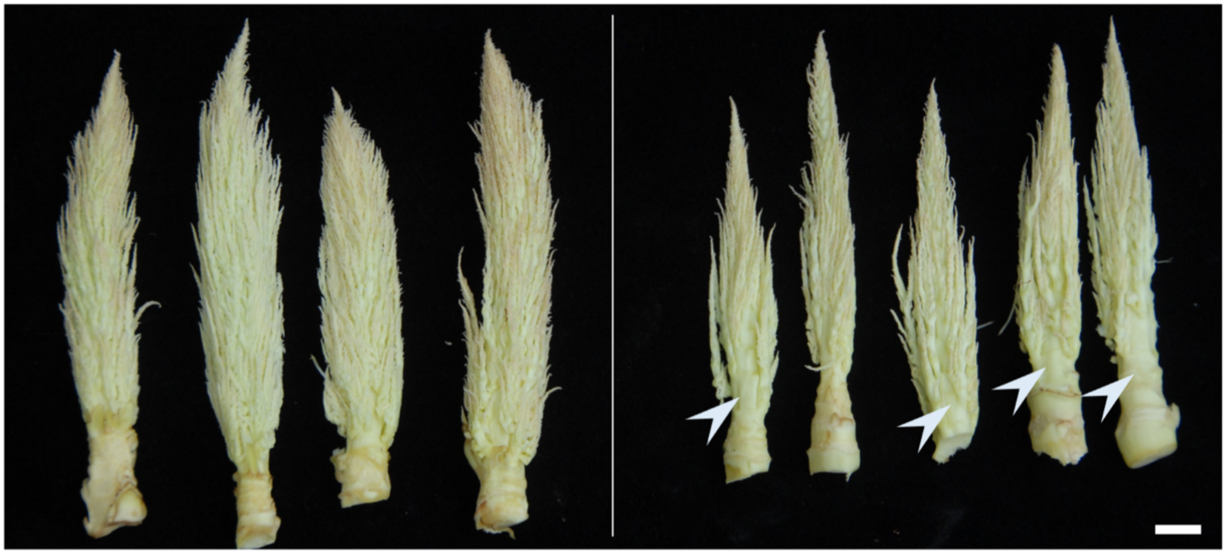

*ra1-R;ra2-R* *ra1-R;ra2-R;o1-2995*

**Supplementary Figure 1.** Ears from a field-grown population segregating double and triple mutant plants. The *ra1-R*;*ra2-R* double mutant ears (left) are sterile and proliferatively branched due to a synergistic interaction that eliminates the majority of *ramosa* pathway function (Vollbrecht et al., 2006). The *ra1-R*;*ra2-R;o1-2995* triple mutant ears (right) are also sterile and synergistically branched, but with overall reductions in growth and size including many fewer branches and relatively large barren, unbranched patches on the ear axis (arrowheads). Bar, 1 cm.

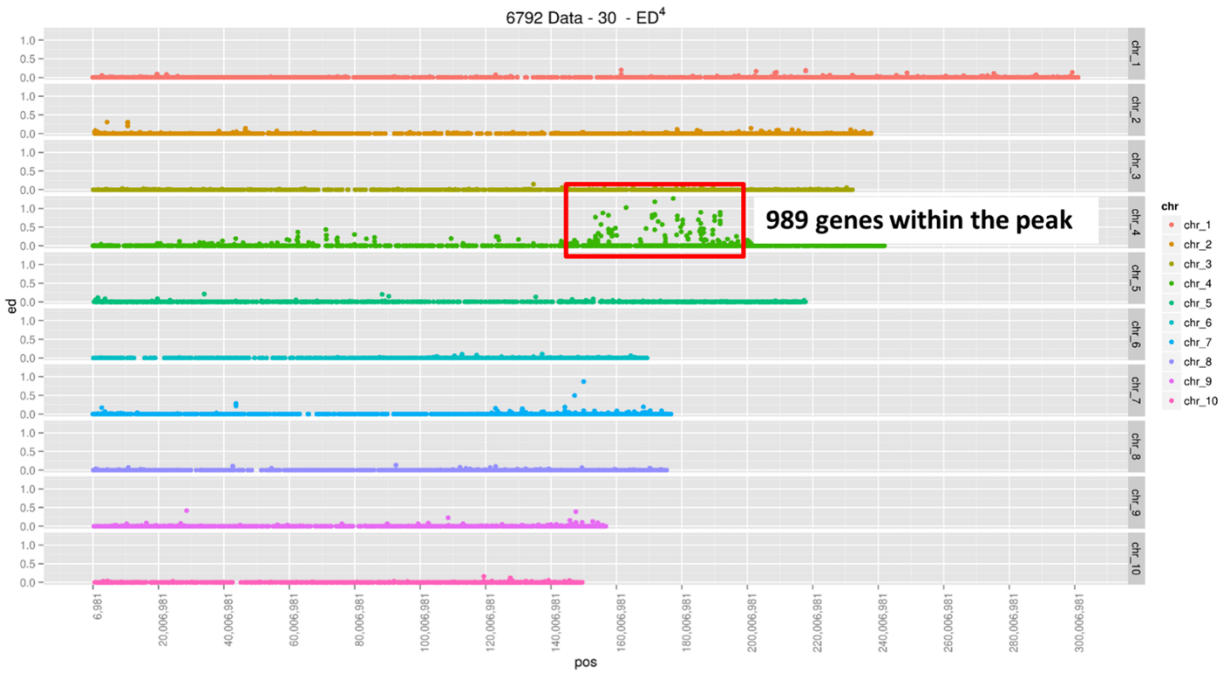


**Supplementary Figure 2.** BSA-GBS rough mapping localizes the *rsl*-12.2995* locus to chromosome 4. Y-axis, left: ed, (Euclidean distance)$^4$ normalized to 1. X-axis: pos, position in bp in the maize B73 RefGen_AGPv4 genome assembly across each of the 10 maize chromosomes (Y-axis, right). Red box, mapping peak on chromosome 4.

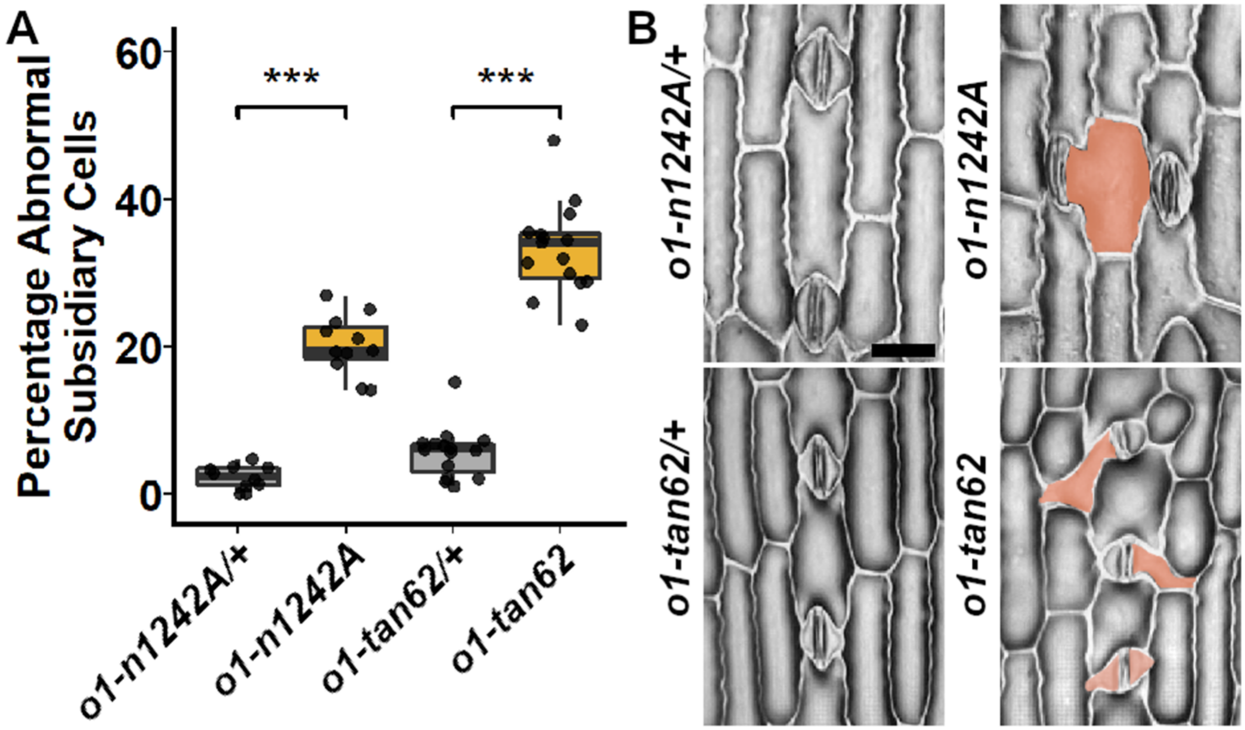


**Supplementary Figure 3.** *o1-tan62* mutants have abnormal subsidiary cell division. **(A)**. Abnormal subsidiary cells were counted from leaf 2 in *o1-tan62* (n=15 plants), *o1-N1242A* (n=11 plants), and their corresponding wildtype siblings (n=15 and 10 plants, respectively). Asterisks (***) indicate significant difference by a two-tailed T-Test, $P < 0.001$. **(B)** Methacrylate impressions on leaf 2 of *o1-n1242A*, *o1-tan62*, and their corresponding wildtype siblings. Abnormal subsidiary cells are highlighted in brown. All images are same magnification, scale bar, 20 μm.

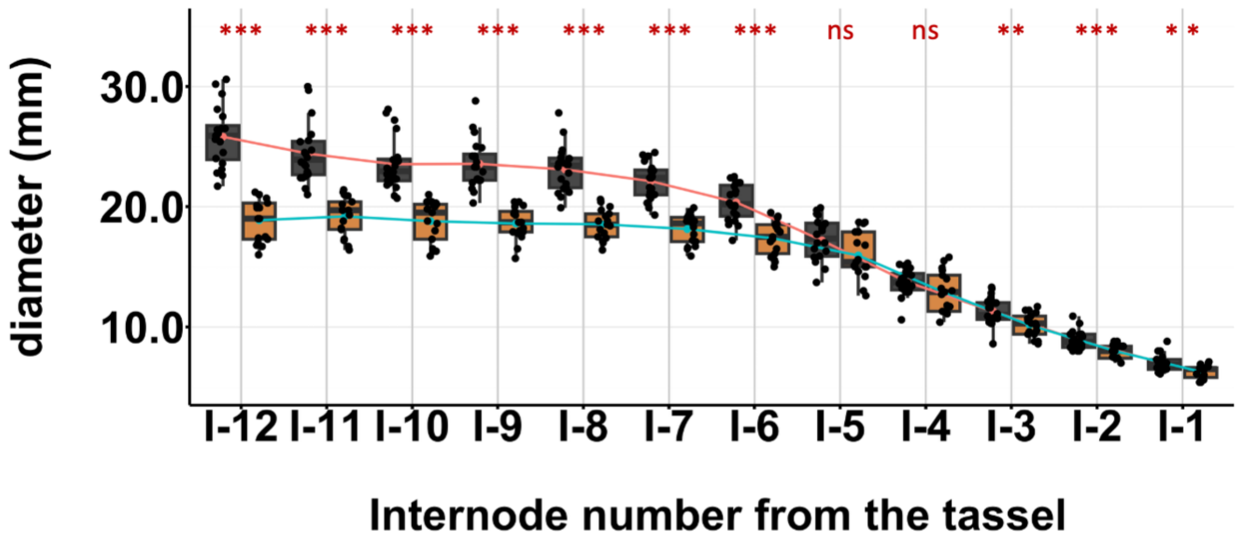


**Supplementary Figure 4.** Internode diameter is compressed in *o1-2995* mutants. Diameter of several internodes of the main shoot in normal siblings (WT, black, left, successive means connected by red line, n = 20) and *o1-2995* mutants (orange, right, successive means connected by blue line, n = 17) in B73. I-1 denotes the internode nearest the tassel and I-12 is 11 internodes away, near the soil line. Asterisks indicate significant difference by a two-tailed T-Test, (*) $P<0.05$; (**) $P<0.01$; (***) $P<0.001$; ns, no significant difference.

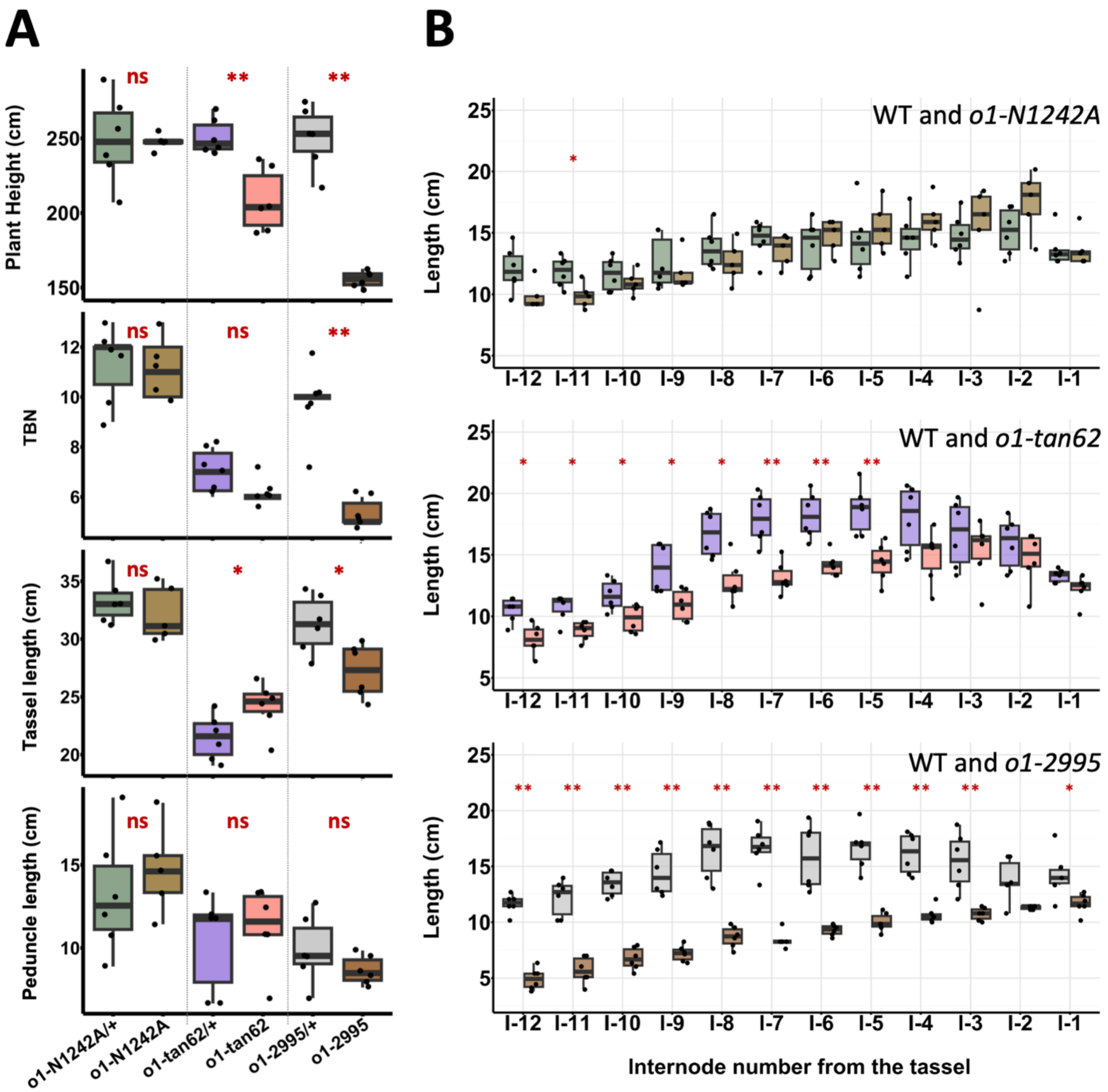


**Supplementary Figure 5.** Phenotypic measurements of different *opaque* mutant alleles in the greenhouse at the University of California Riverside. For all measurements: *o1-N1242A* (n=5 plants), *o1-tan62* (n=6 plants), *o1-2995* (n=6 plants) and their respective wild type (WT) siblings (n=6 plants each). **(A).** *o1-N1242A/+* siblings and *o1-N1242A* null mutants (left); *o1-tan62/+* siblings and *o1-tan62* mutants (center); *o1-2995/+* siblings and *o1-2995* antimorph mutants (right). Traits measured are Plant height (cm); TBN, tassel branch number; Tassel length (cm); Peduncle length (cm). **(B).** Boxplots indicating internode length (cm) for each internode from nearest the tassel (I-1) to near the soil (I-12) for different *opaque* mutants and their wild type (WT) siblings. Asterisks indicate significant difference by a two-tailed T-Test, (*) $P<0.05$; (**) $P<0.01$; ns or no asterisk, no significant difference.

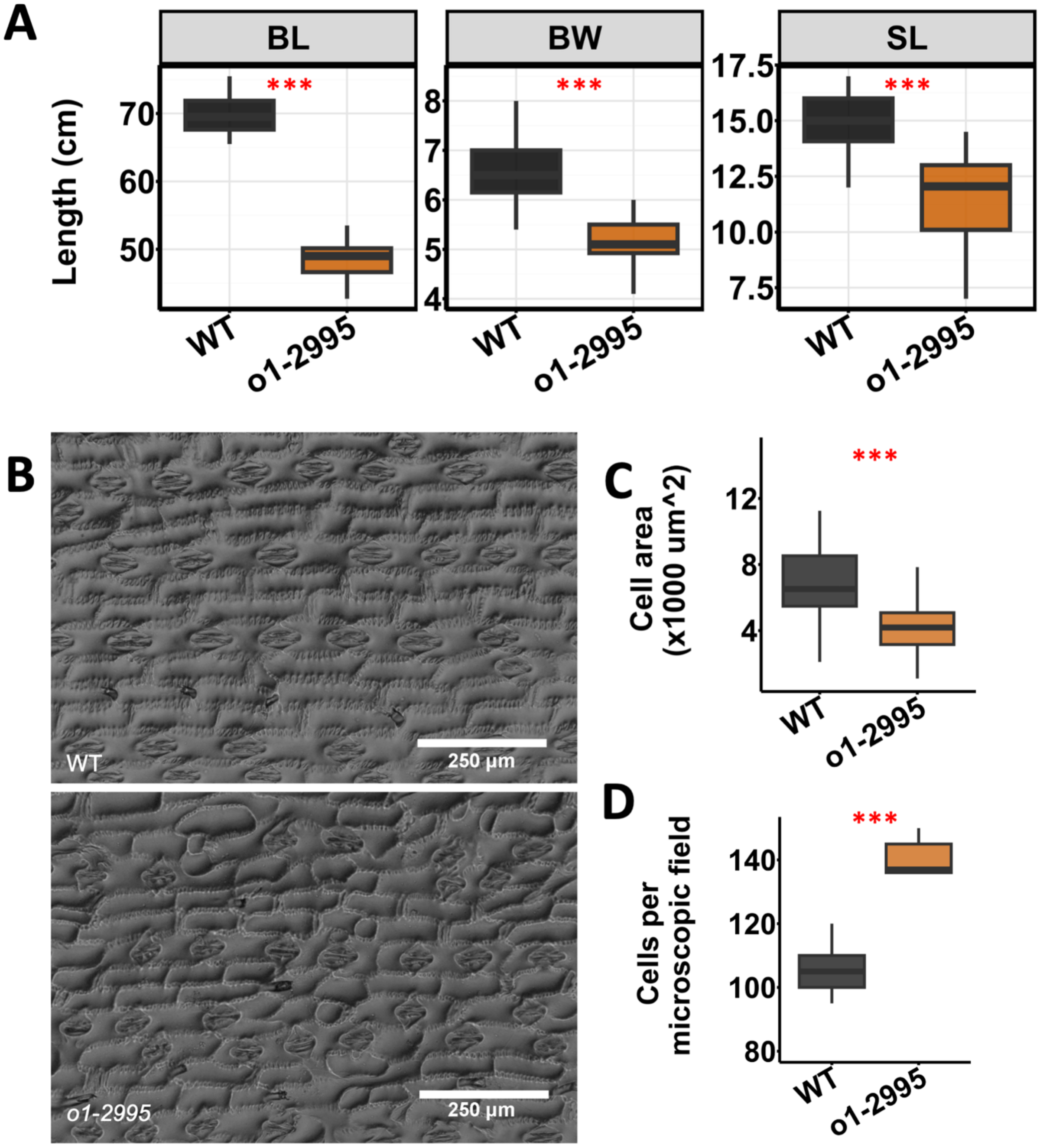


**Supplementary Figure 6.** *o1-2995* mutation reduces leaf size and affects subsidiary cell division. **(A).** Leaf traits measured for leaf 16 normal (n=10) and sibling *o1-2995* (n=10) plants. BL, blade length; BW, blade width; SL, sheath length. **(B)**. Representative leaf 16 epidermal impression of the wild type siblings (WT, upper) and *o1-2995* mutants (lower). **(D)**. Quantification of cell parameters from bright field microscope images of leaf 16 epidermal impression of the wild type siblings (WT, n=5) and *o1-2995* mutants (n=5). **(C)**. Quantification of cell area (*** P = 0.0012). **(D).** Quantification of cell number per microscope field (*** P = 1.5e-09).

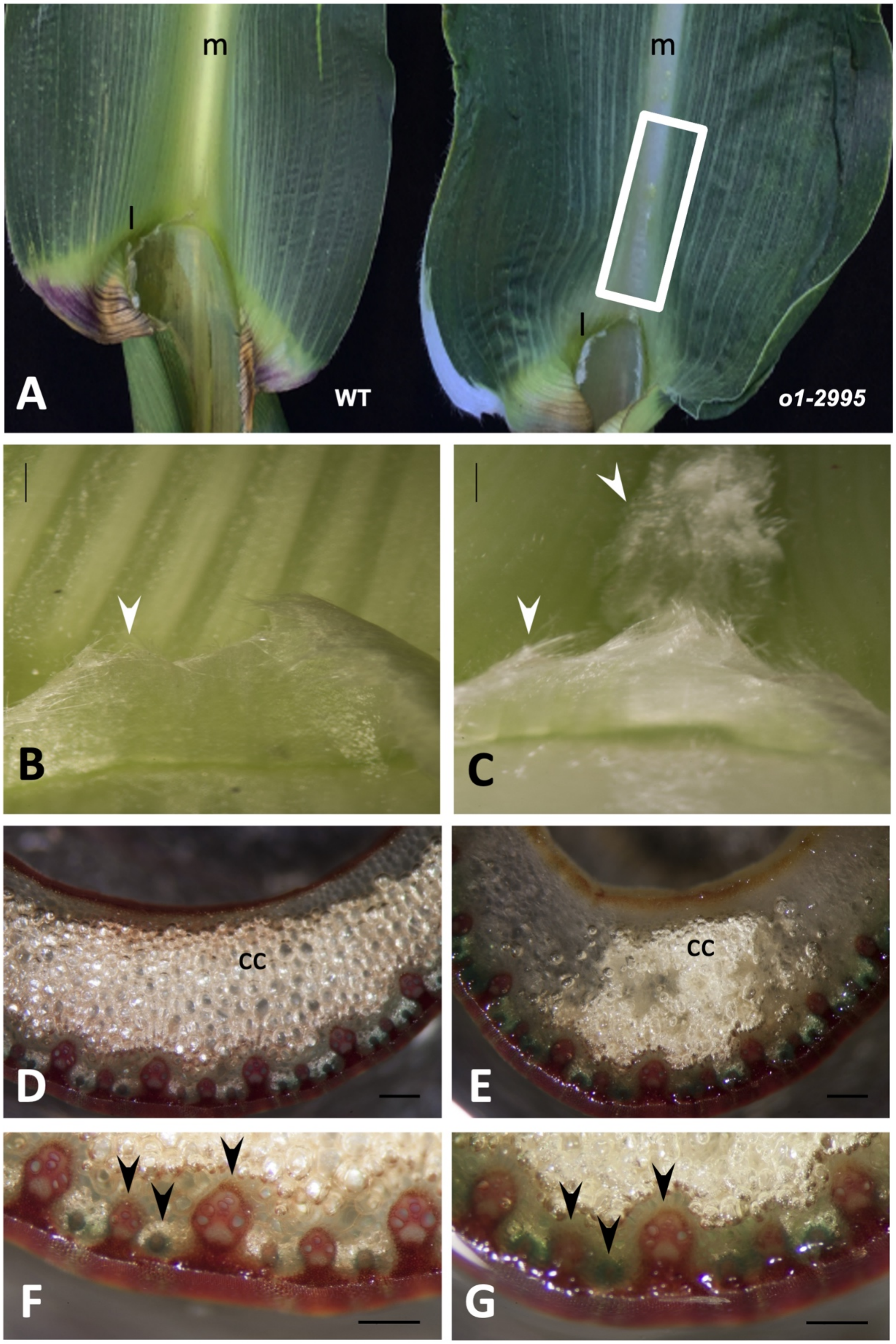


**Supplementary Figure 7:** The *o1-2995* mutation impacts leaf patterning: **(A to C)**. *o1-2995* mutants have altered leaf patterning. m, midrib; l, ligule. **(A)**. *o1-2995* mutant (right) with ectopic tissue out-growths across the leaf midrib (area within white box) compared a normal sibling (WT, left). **(B and C)**. Close-up of ligule fringe (white arrowheads) at the midrib on upper (adaxial) leaf surface. **(B)**. Normal sibling. **(C)**. *o1.2995* mutant with ectopic ligule located distally, in the midrib. (**D-G**). Leaf midrib staining with acidified phloroglucinol. Cross sections of midrib of adult leaf 10, taken 10 cm distal to the ligule. CC, clear cells. **(D)** Normal sibling, with a portion of the midrib in the field of view. Black arrowheads, vascular bundles. **(E)** *o1-2995* mutant, same magnification as (D), with the entire midvein in the field of view. The midvein is reduced in *o1-2995* mutants compared to in normal siblings, as indicated by the relatively reduced area of clear cells. **(F)**. Normal sibling; higher magnification image of abaxial midrib section shown in (D). **(G)**. *o1-2995* mutant; higher magnification image of abaxial midrib section shown in (E); in *o1-2995* mutants the vascular bundles (e.g., arrowheads) are qualitatively smaller than analagous-order veins in WT siblings in (F). Bars: B and C, 400 µm; D and E, 400 µm; F and G, 250 µm.
